# The Role of Evolving Interfacial Substrate Properties on Heterogeneous Cellulose Hydrolysis Kinetics

**DOI:** 10.1101/691071

**Authors:** Jennifer Nill, Tina Jeoh

**Author notes:** Corresponding author: Tina Jeoh.

## Abstract

Interfacial enzyme reactions require formation of an enzyme-substrate complex at the surface of a heterogeneous substrate, but often multiple modes of enzyme binding and types of binding sites complicate analysis of their kinetics. Excess of heterogeneous substrate is often used as a justification to model the substrate as unchanging; but using the study of the enzymatic hydrolysis of insoluble cellulose as an example, we argue that reaction rates are dependent on evolving substrate interfacial properties. We hypothesize that the relative abundance of binding sites on cellulose where hydrolysis can occur (productive binding sites) and binding sites where hydrolysis cannot be initiated or is inhibited (non-productive binding sites) contribute to rate limitations. We show that the initial total number of productive binding sites (the productive binding capacity) determines the magnitude of the initial burst phase of cellulose hydrolysis, while productive binding site depletion explains overall hydrolysis kinetics. Furthermore, we show that irreversibly bound surface enzymes contribute to the depletion of productive binding sites. Our model shows that increasing the ratio of productive- to non-productive binding sites promotes hydrolysis, while maintaining an elevated productive binding capacity throughout conversion is key to preventing hydrolysis slowdown.

## Introduction

Interfacial enzyme reactions, which are enzymatic reactions that occur at interfaces, including membrane bound and other surface immobilized enzymes^1^, lipases which act at lipid-water interfaces^2^, and enzymes which bind to and degrade insoluble fibers at the solid-liquid interface such as collagenases^3^, amylases^4^ and cellulases^5^. These reactions are widespread *in vivo* and industrially, but the underlying rate governing mechanisms of these reactions are often obscured by complex interplay of reaction intermediates and reaction steps. For example, the enzymatic hydrolysis of insoluble cellulose has long been studied, yet full mechanistic understanding of the reaction of cellulose degrading enzymes called cellulases, which produce soluble sugars from cellulose, remains elusive. An archetypal product release curve for hydrolysis of cellulose by cellulases is shown in Figure 1, where an initially rapid period of product release (the burst phase) is followed by a decay of product release rates during the slowdown and plateau phases, despite the remaining presence of cellulose in the reaction. Product inhibition and enzyme instability only partially account for this slowdown^6^ and the underlying mechanisms causing rate slowdown have been probed with numerous kinetic models^7, 8^, yet the molecular origins of the rate slowdown of the enzymatic hydrolysis of cellulose remains incomplete.

**Figure 1:**
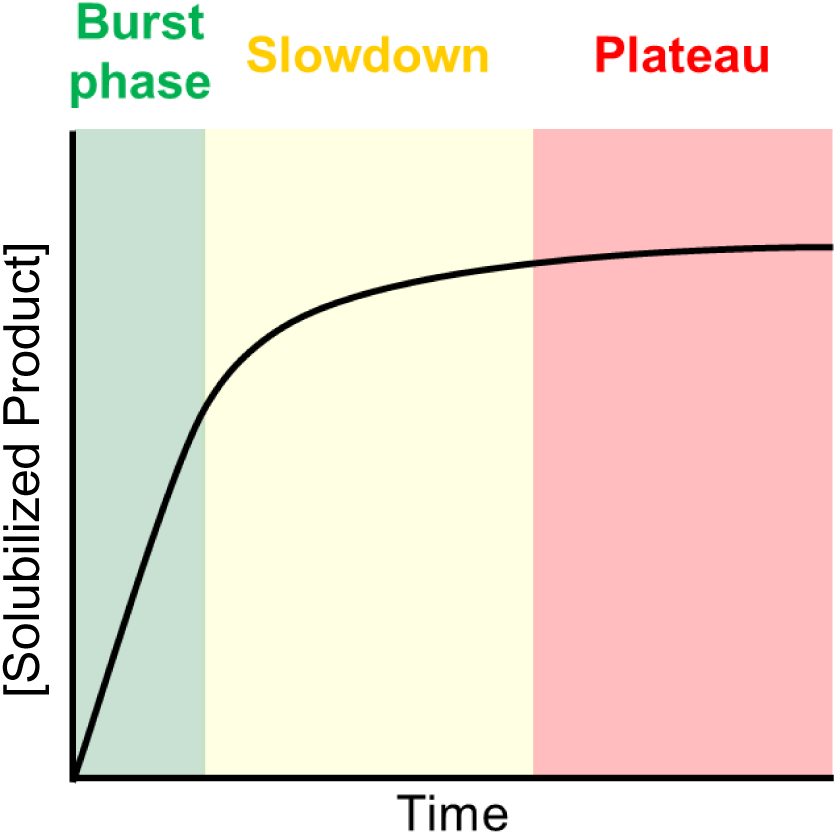
Representative cellulose hydrolysis curve depicting solubilized product concentration as a function of time. Cellulose hydrolysis is characterized by a burst phase, where maximum hydrolysis rates are reached early in the reaction, and progressive slowing of product release with eventual decay to negligible release of product, despite reaching incomplete conversion.

In order for an exocellulase to successfully hydrolyze cellulose, it must first adsorb to the insoluble surface of cellulose and thread a single chain of cellulose in its catalytic domain in a process termed complexation, before it processively catalyzes multiple cleavages of soluble sugars from the cellulose particle (Figure 2). Studies elucidating inherent enzyme properties contributing to rate limitations have pointed to the processivity^9–12^, on-rate^13–15^, or off -rate^10, 12, 16–19^ as inherent properties of cellulases that lead to hydrolysis slowdown, but these models often fail to predict hydrolysis trends in their entirety^8^. Understanding of enzyme binding to cellulose is essential for full elucidation of hydrolysis kinetics, but the two domain structure of exocellulases like *Trichoderma reesei* Cel7A allows for multiple modes of enzyme binding by the carbohydrate binding module (CBM) and the active-site-containing catalytic domain (CD)^19–20^. An enzyme that is complexed with a cellulose chain and producing soluble product is said to be *productively bound*, and we define the total number of sites per mass of cellulosic substrate wherein productive binding can occur as the *productive binding capacity*. Conversely, enzyme bound to the surface in the inactive state (*non-productively bound*) can be non-specific, by the CBM only, or complexed by the CD but somehow stalled or inactivated^10, 13, 14, 20, 21^, as illustrated in Figure 2. Experimental quantification of bound enzyme with occupied or free active sites has been performed using a soluble reporter molecule^9, 17, 22–25^, but this method does not differentiate between productively bound enzymes and catalytically inactive enzymes with an occupied active site.

**Figure 2:**
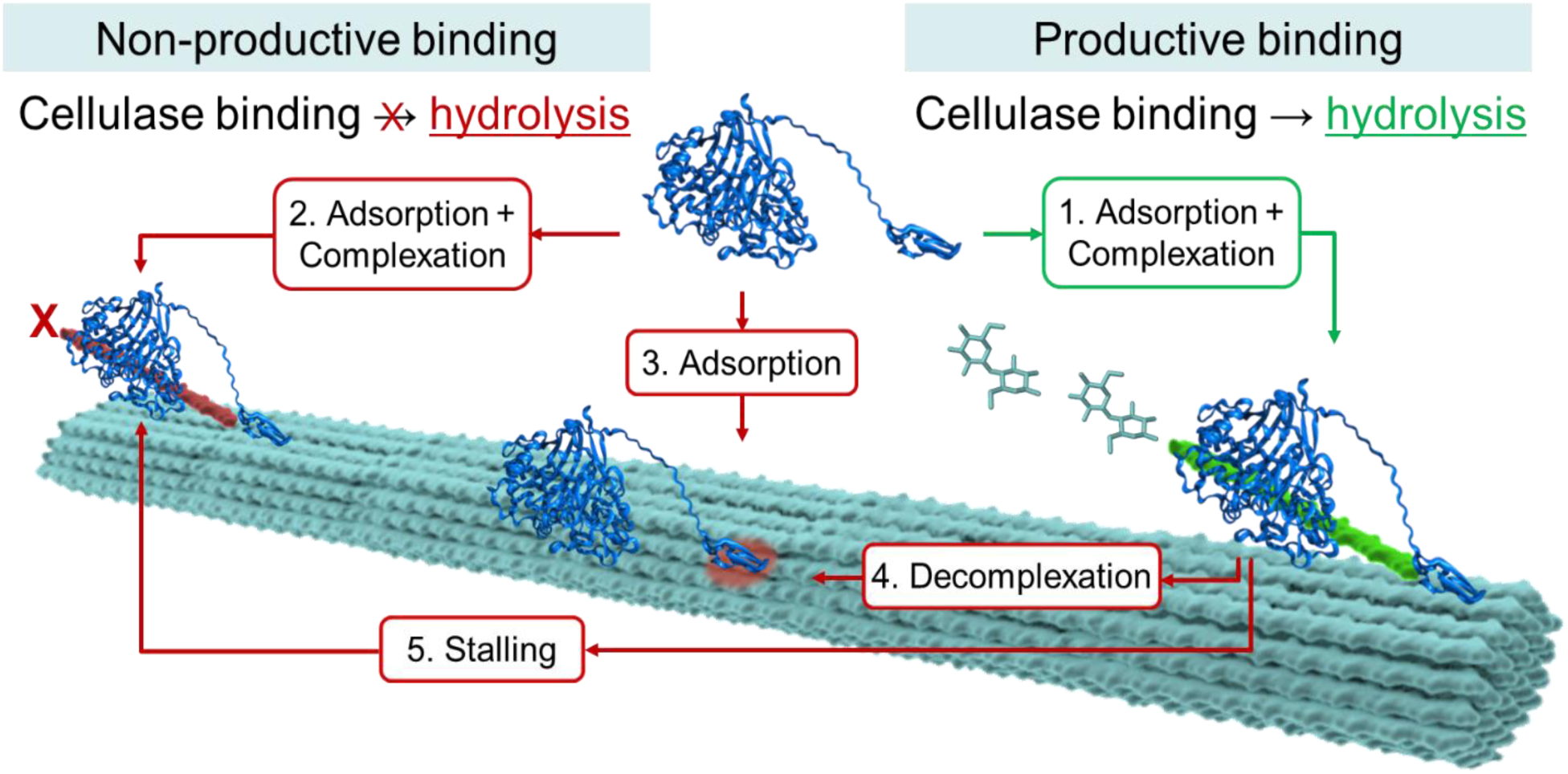
Illustration of binding modes of Cel7A to a cellulose fibril. Productive binding occurs when a Cel7A molecule (1) adsorbs and complexes with an accessible cellulose chain (a productive binding site) resulting in hydrolysis. Non-productive binding can occur by the following mechanisms: (2) enzyme in solution adsorbs and complexes with a non-hydrolyzable chain end, (3) enzyme in solution adsorbs by the CBM only, (4) a productively bound enzyme decomplexes to become bound by the CBM only, or (5) a productively bound enzyme becomes blocked on the surface when it encounters a surface obstacle (a non-productive binding site or another non-productively bound enzyme) in a process referred to as ‘stalling’. Non-hydrolyzable chain ends, surface obstacles causing stalling, or adsorption binding sites are lumped into the category of ‘non-productive binding sites’. Enzyme was constructed in PyMol^15^ from (PDB 4C4C)^16^, (PDB 1CBH)^17^, and using the linker sequence from Badino et al. 201718.

Further complicating our understanding of hydrolysis kinetics is the substantial effect of cellulose morphology on hydrolysis rates, and many studies conclude that rate loss is substrate limited^8, 15, 26–31^. However, understanding of the role of inherent cellulose properties and enzymatic modification of the substrate in cellulose hydrolysis remains incomplete, with limited understanding of substrate properties affecting hydrolysis both initially and longer term. For instance, surface obstacles are often cited as an intrinsic substrate property that causes premature termination of cellulolytic action9, 10, 13, 15, 20-22, 26, 32-35, but their structural origin remains obscure. Cellulose accessibility is described as another key parameter which limits hydrolysis, and a number of previous reports have estimated the concentration of productive binding sites on a variety of celluloses^17, 22, 23, 31^, but understanding of what makes a cellulose chain accessible to hydrolysis by cellulases is limited. Studies often point to the crystallinity of cellulose as the origin of cellulose reacalcitrance^27, 36^, as the free energy required to form a productive complex from a cellulose crystal is substantial^37, 38^, but it was recently shown that the initial accessibility of cellulose is not governed exclusively by crystallinity.^31^ The kinetics of cellulose hydrolysis are also highly dependent on solids loadings and enzyme-to-substrate ratios, and the apparent rate limiting factors can vary greatly depending on the chosen reaction conditions.^11, 39, 40^

The effect of enzyme action on substrate properties limiting hydrolysis is largely uncharacterized. Often, attempts to understand the role of cellulose heterogeneity in rate decrease involves descriptions of more- and less-accessible fractions of substrate^41, 42^, but understanding of the evolution of the accessible fraction—i.e., the productive binding capacity—throughout hydrolysis is limited. As the heterogeneous reaction takes place at the solid-liquid interface, it is reasonable to assume a surface ablation mechanism by which removal of productive binding sites from an accessible fraction exposes new sites from an inaccessible central core ^13, 17, 32, 39, 40, 43–50^, and the initial cellulose surface area and concentration of productive binding sites has been shown to limit hydrolysis rates^8, 11^, in support of this mechanism. However, mechanically increasing cellulose chain reducing end concentrations did not improve cellulose accessibility to *Tr*Cel7A (a reducing-end specific cellobiohydrolase)^48^, and the availability of reactive sites is not proportional to surface area or total reducing end concentrations^31, 51^, indicating that the location of a cellulose chain on the surface of a cellulose particle alone is not a sufficient criterion to indicate accessibility. Furthermore, neither depletion of surface area nor shortening of chains explains long term hydrolysis slowdown^8^. Previous studies have linked structural effects of enzyme degradation to accessibility^41, 42, 48, 52–55^, but to our knowledge, the results shown here are the first specifically quantifying the accessibility of cellulose by means of productive binding capacity measurements throughout the duration of hydrolysis.

Since the true physical definition of a productive binding site remains undefined, direct experimental measurement of the concentration of productive binding sites (sometimes also referred to as attack sites) of a cellulosic substrate has not been reported. It is, however, possible to use the enzyme as a reporter to estimate the concentration of productive binding sites. Using the ratio of specific maximum rates obtained at enzyme and substrate saturation, the concentration of productive binding sites has been estimated on Avicel^40^. Here, we use the rate of product released by Cel7A as a reporter of the concentration of binding sites, which at saturation directly correlates to the number of available productive Cel7A binding sites on cellulose, shown schematically in Figure 3.

**Figure 3:**
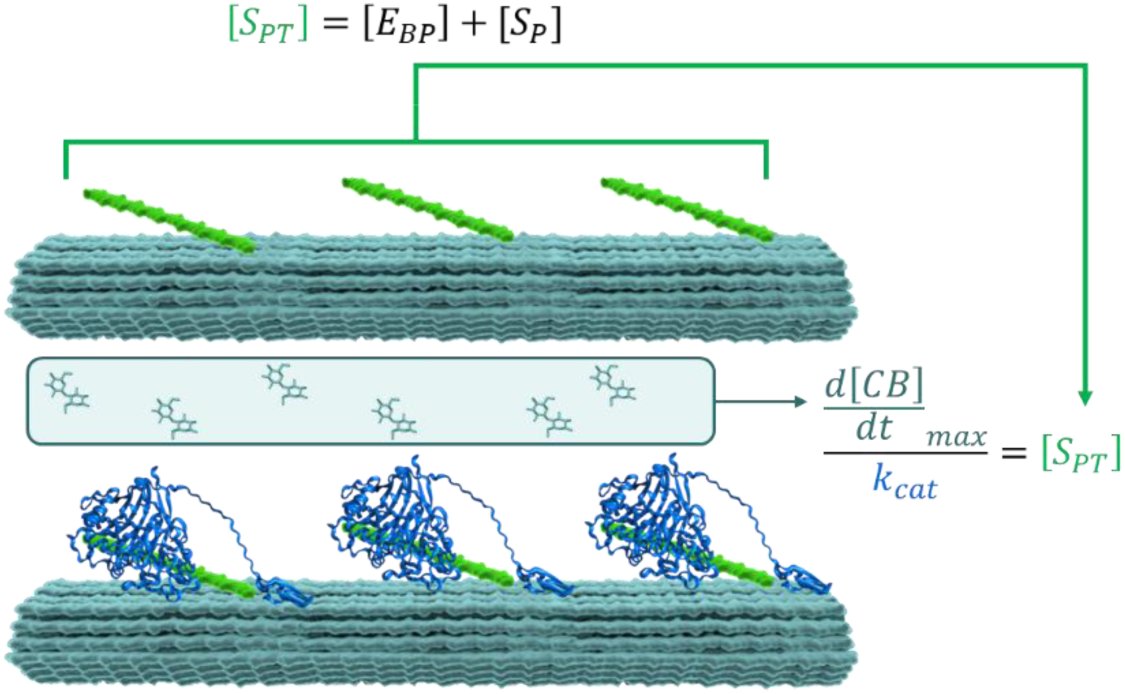
Schematic illustration of the estimation of the productive Cel7A binding capacity ([S_PT_], µmoles/g) of cellulose by saturating the available productive binding sites with Cel7A and measuring cellobiose (CB) release rates, effectively ‘counting’ the number of productive binding sites. The productive Cel7A binding capacity is the sum of productive binding sites that are unoccupied ([S_P_]) and occupied by productively bound enzyme ([E_BP_]); it is a measure of the maximum number of available productive binding sites per mass of cellulose.

We previously simulated the kinetics of cellulose hydrolysis by Cel7A using a mechanistic model based on the interactions described in Figure 2. When the concentration of productive binding sites was held constant at the initial productive binding capacity of the substrate ([S_P_]_0_ = [S_PT_] and d[S_P_]/dt=0), the model accurately captured burst phase product release rates, but over-predicted beyond the first several seconds of the reaction^6, 26^. In the current work, we quantified the time dependent change in the productive Cel7A binding capacity of cellulose. Incorporating these trends into our model allowed us to accurately predict hydrolysis trends for a number of substrates, and provides important insight into the true origins of cellulose recalcitrance.

## Experimental Section

### Purification of Cel7A

Cel7A was purified from a commercial cellulase preparation (Sigma Cat# C2730) by anion exchange, *p*-aminophenyl-β-d-cellobioside (*p*APC) affinity, and size exclusion chromatography as described previously^53^ with the addition of a buffer exchange into 50 mM sodium acetate, 100 mM sodium chloride (pH 5.0) preceding loading onto the size exclusion column. The *p*APC column matrix was synthesized using commercially available *p*APC (Carbosynth). Purity of the final Cel7A preparation was identified by a single band in an SDS-PAGE at ∼65 kDA and the identity of Cel7A was verified by LC/MS/MS at the proteomics facility at University of California, Davis (http://proteomics.ucdavis.edu/).

### Cellulose Preparation

Microcrystalline cellulose (MCC) (Acros Organics, cat#382312500) and Filter paper (FP) (Whatman #1, Piscataway, NJ) were suspended in 50 mM sodium acetate and blended in a Waring blender. Swollen Filter Paper (SC) was prepared from filter paper by incubating overnight in cold phosphoric acid (85% w/v) and regenerating in cold water.^31^ Bacterial Cellulose from *Gluconacetobacter xylinum* (BC) cultured in static cultures and algal cellulose from *Cladophora aegagropila* (AC) were prepared as previously described^31, 56^. Hydrochloric acid treated algal cellulose (HCl AC) was prepared with adjustments to previously published methods^57^. Algal cellulose was incubated with 5 M HCl at 70 °C overnight and then extensively washed with water and 50 mM sodium acetate. Dried cellulose was generated by drying never-dried SC, BC and AC at room temperature and resuspending in buffer.

### Preparation of partially hydrolyzed cellulose

Cellulose that was hydrolyzed to varying extents by Cel7A was generated and washed using modifications to methods by Jung et al.^58^, Jeoh et al.^59^ and Yang et al.^21^. Saccharification of FP, SC, MCC, BC, and AC was conducted using 2.5 µM or 0.25 µM purified Cel7A with 0.5 mg/mL substrate loading in 50 mM sodium acetate (pH 5.0) in 5 mL CENTREX 0.45 um Nylon filter tubes. The filter tube bottoms were plugged and the reactions were conducted at 50 °C with 25 rpm end over end rotation. The reactions were stopped after 0, 1, 4, 24, 48 and 120 hours for FP, AC, and MCC, and 0, 5, 15, 45 and 60 minutes for BC and SC by centrifuging the tubes at 4000 rpm and 4 °C for 1 minute. The cellulose was immediately washed to remove residual active enzyme with either a salt wash or protease wash as described below. The error in solids loading due to losses during washing was determined to be negligible by an Anthrone assay before and after washing and is detailed in Supporting Information (SI). The washed substrates were analyzed with Synchrotron FTIR (sFTIR), using the presence or absence of the amide I and II peaks between 1500 and 1700 cm^-1^ as indication of the presence or absence of surface bound Cel7A (SI). Extent of hydrolysis was calculated from the concentration of soluble sugars in the filtrate measured as described below.

### Cellulase removal from partially hydrolyzed cellulose

Partially hydrolyzed cellulose retentates were resuspended in 0.5 M NaCl and filtered. This process was repeated twice. The cellulose was then washed in a similar manner using 50 mM sodium acetate buffer pH 5.0 three times and resuspended. To remove any irreversibly bound protein, a portion of the salt washed cellulose was resuspended in phosphate buffer (pH 7.5) and incubated with 0.2 mg/mL Pronase E at 25 rpm and 37 C overnight.^21^ The protease was then washed from the cellulose using the same salt washing protocol as described above.

### Quantification of solubilized sugars

The soluble sugars in the filtrate were measured by high performance anion exchange chromatography with pulsed amperometric detection (HPAE-PAD) using Dionex™ CarboPac™ SA10 Analytical and guard columns (Dionex, Thermo Fisher Scientific Inc., Sunnyvale, CA) as described previously^31^.

### Determining the productive Cel7A binding capacity of cellulosic substrates

The productive binding capacity was estimated from maximum hydrolysis rates from monitoring the concentration of cellobiose (CB) released over short times (d[CB]/dt) within the burst phase of hydrolysis as previously described^11, 31^. Enzyme loadings ranging from 5-130 µmoles Cel7A/g of cellulose were added to 0.05 mg/mL of cellulose in a stirred, jacketed cell at 50°C and 200 rpm to initiate each reaction. An amperometric cellobiose biosensor^60^ using *Phanerochaete chrysosporium* cellobiose dehydrogenase (CDH) was used to obtain real-time (sub-second) measurements of cellobiose concentrations within the first minute of the reaction at a sampling rate of 10 s^-1^. The productive Cel7A binding capacities of SC, BC, FP, MCC and AC were previously measured by Karuna and Jeoh from fits to discretely sampled cellobiose concentrations at time points within a few seconds at the start of each hydrolysis reaction.^6^ In this study, the use of a CDH biosensor significantly improved this measurement with real-time, continuous measurements of cellobiose release in the hydrolysis reaction. A comparison of the productive binding capacities estimated for the five celluloses show good agreement and a marked increase in experimental reproducibility (Figure 4). Details of the biosensor, including calibration, validation and example data are provided in supporting information.

**Figure 4:**
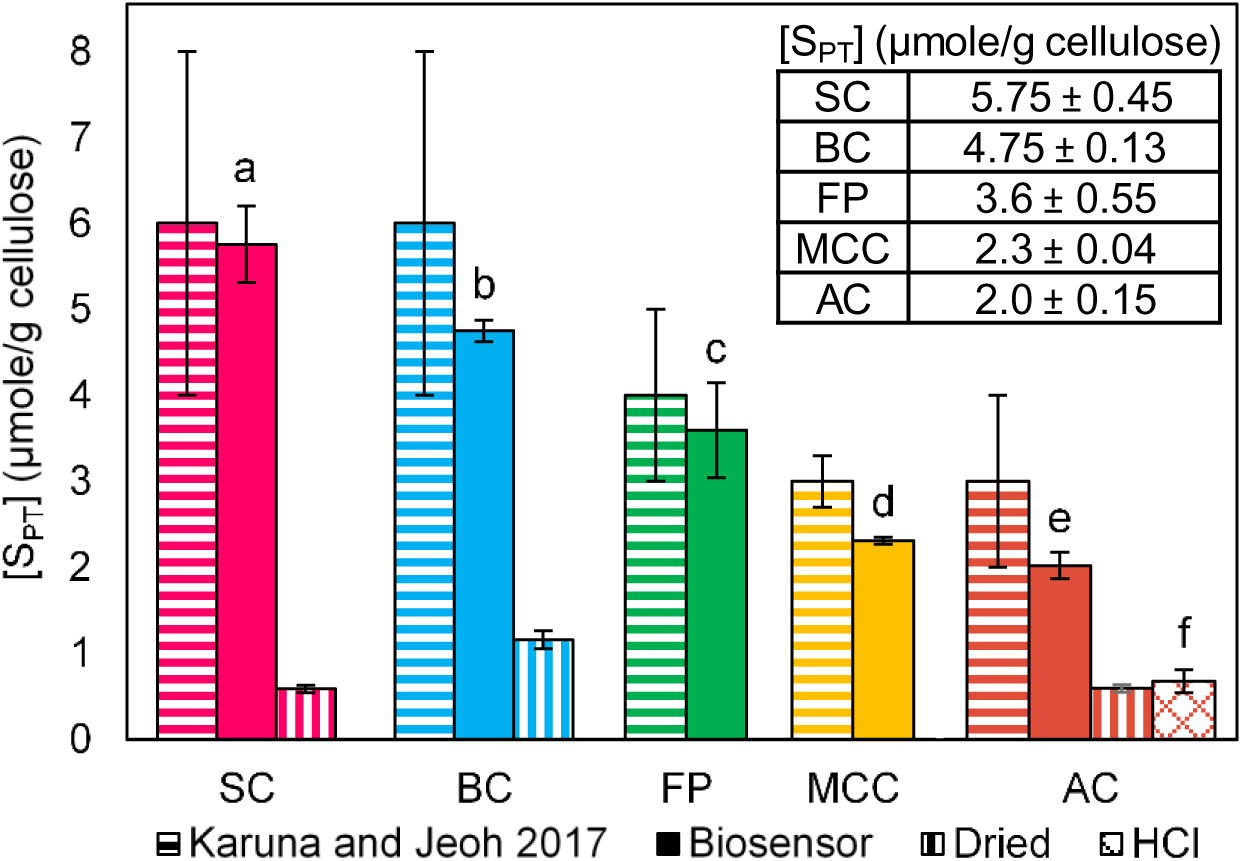
The productive binding capacities of filter paper (green bars), microcrystalline cellulose (yellow bars), algal cellulose (red bars), bacterial cellulose (blue bars) and swollen cellulose (pink bars). Results from cellobiose release rate measurements by discrete sampling (speckled bars) and real-time, continuous sampling using an amperometric CDH biosensor (solid bars) are shown to be comparable. The productive binding capacity of dried cellulose (diagonal striped bars) and HCl hydrolyzed cellulose (crosshatched bars) were also measured using a biosensor. Unique lowercase letters above bars indicate a statistically significant difference between means (α = 0.05), as determined by a 2-sided t-test between all means shown for the productive binding capacity of never dried cellulose measured using a CDH biosensor. The experimental means and error from triplicate measurements (n=3) for each cellulose are tabulated in the inset.

The concentration of productively bound Cel7A ([E_BP_]) was estimated from the first order relationship (Equation 1) describing processive hydrolysis of cellulose by a productively bound enzyme with k_cat_ = 5.4 s^-1^.^11^

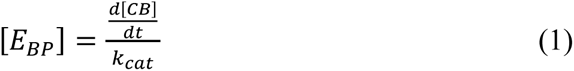

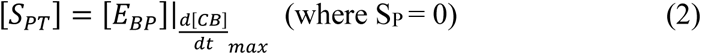

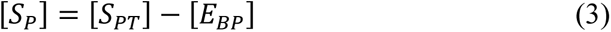

The productive binding capacity, S_PT_ was estimated as the concentration of productive binding sites at saturating enzyme loadings (i.e., no further increase in initial hydrolysis rates with additional enzyme loading) (Figure 3 and Equations 2-3).

### Estimating the time dependency of the productive binding capacity of cellulose during hydrolysis

The productive binding capacities as a function of time and conversion for all celluloses were fit using the method of Maximum likelihood to minimize the Chi-squared (χ^2^) statistic. Using the Levenberg-Marquadt algorithm to minimize χ^2^, we fit the data to one- and two-exponential decay functions and selected fits with a reduced χ^2^ value (χ^2^/Degrees of Freedom) closest to 1.^61^ A single-exponential decay equation under-fit the data in Figure 5 (χ^2^_reduced_ >> 1), while a double-exponential decay (Equation 4) fit to the data resulted in χ^2^_reduced_ < 1 (Tables 3 and S1). Thus the productive binding capacities for all celluloses with respect to time were best fit by a double exponential decay equation, suggesting that there are two populations of productive binding sites (Equation 4):

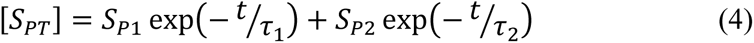

where S_PT_ is the total concentration of productive binding sites (µmole/g), S_P1_ and S_P2_ are pre-exponential factors describing initial concentrations of accessible sites (µmole/g), and τ_1_ and τ_2_ are characteristic decay times of the two productive binding site populations (h). Equation 4 was differentiated with respect to time to empirically model the rate of change of productive binding sites on cellulose during hydrolysis:

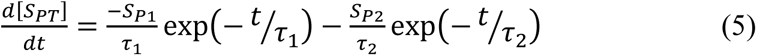

**Figure 5:**
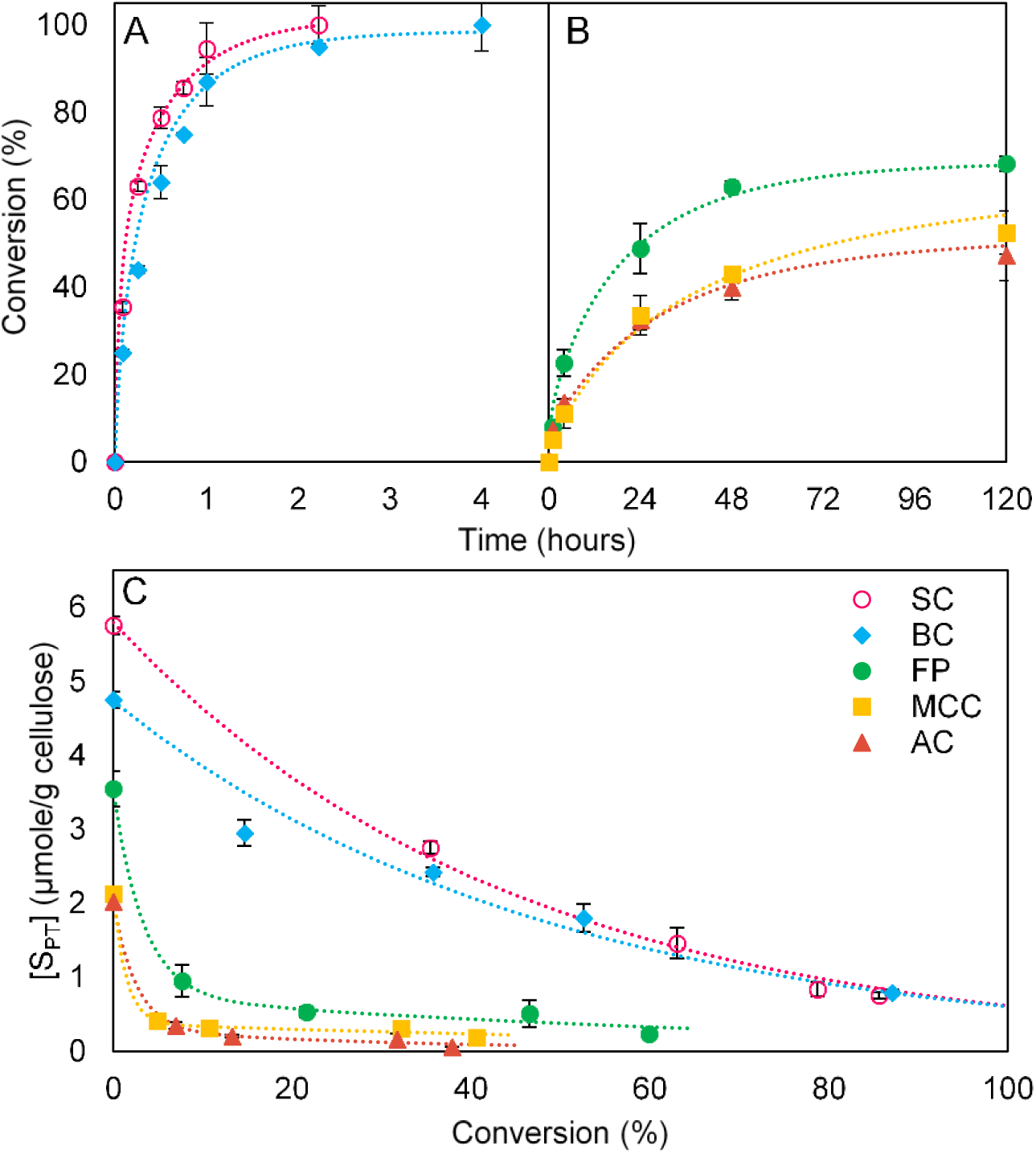
A) Hydrolysis curves of swollen cellulose (SC) and bacterial cellulose (BC); B) Hydrolysis curves of algal cellulose (AC), filter paper (FP), and microcrystalline cellulose (MCC); C) The evolution of productive binding capacity of cellulose as a function of conversion. Partially hydrolyzed celluloses were washed extensively with a salt solution prior to the measurement of the productive binding capacity of the ‘salt washed cellulose’ (SWC). Dotted lines in (C) are fits using Equation 4 (Fitting parameters are summarized in Table 3); dotted lines in (A) and (B) are simulation results from incorporating the scheme in Table 1 and Equation 3.

**Table 1:**
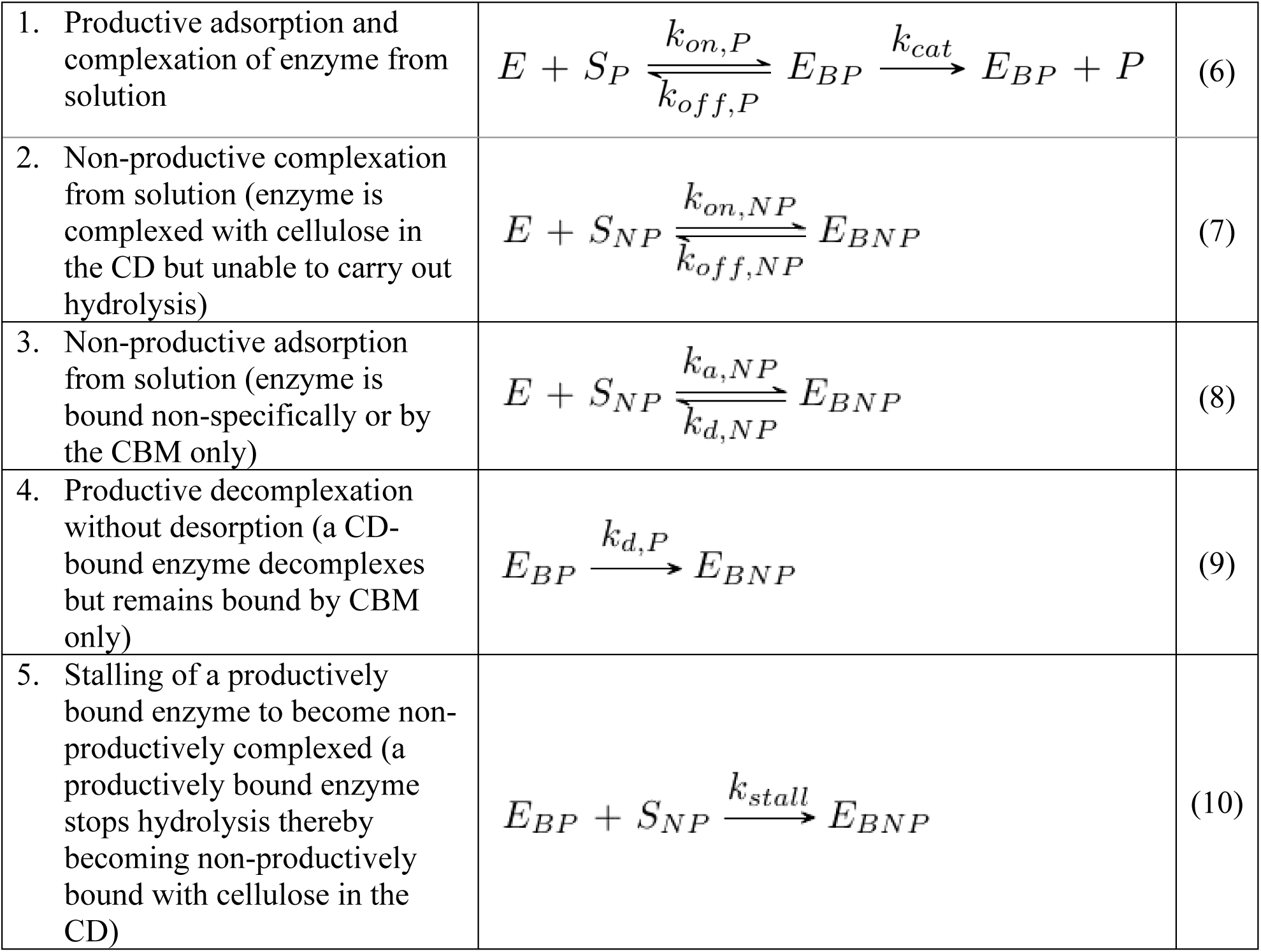
Elementary steps in the overall mechanism of Cel7A hydrolysis of cellulose. Numbering in the scheme corresponds to Figure 2.

### Modeling the hydrolysis of cellulose by Cel7A

The kinetics of cellulose hydrolysis was modeled as described in Table 1 and Figure 2. Equation 6 describes productive interactions between Cel7A and cellulose including: adsorption and complexation of Cel7A enzyme (E) with a productive binding site (S_P_); processive hydrolysis of cellulose by a productively bound enzyme (E_BP_) producing the product (P), cellobiose; and decomplexation/desorption of the enzyme from the insoluble cellulose surface.^11^ Additionally, non-productive binding interactions depicted in Figure 2 where a non-productively bound enzyme (E_BNP_) is formed at non-productive binding site (S_NP_) or from a productively bound enzyme are incorporated. Our model does not differentiate different types of non-productive binding sites, and thus Equations 7-10 describe various mechanisms of non-productive binding, all occurring at S_NP_. While this may be an oversimplification of mechanisms of non-productive binding, the work here focuses on productive binding, and therefore further refinement of the model is outside of the current scope. Elementary steps in Table 1 are numbered according to Figure 2. Descriptions of the rate constants and nomenclature in Table 1 are provided in Table 2.

**Table 2:** Rate constants averaged from the literature (from Nill et al. 2018^11^) and variables used in the simulation of cellulose hydrolysis.

| Rate Constants |  |  |
| --- | --- | --- |
| **k<sub>cat</sub>** | 5.4 | Catalytic rate constant, s<sup>-1</sup> |
| **k<sub>on,P</sub>** | 6.43 x 10<sup>-4</sup> | Productive on-rate, g/μmole/s |
| **k<sub>off,P</sub>** | 0.038 | Productive off-rate, s<sup>-1</sup> |
| **k<sub>a,NP</sub>** | 6.43 x 10<sup>-5</sup> | Non-productive adsorption rate, g/μmole/s |
| **k<sub>d,NP</sub>** | 0.347 | Non-productive desorption rate, s<sup>-1</sup> |
| **k<sub>on,NP</sub>** | 6.43 x 10<sup>-4</sup> | Non-productive on-rate, g/μmole/s |
| **k<sub>off,NP</sub>** | 0.038 | Non-productive off-rate, s<sup>-1</sup> |
| **k<sub>d,P</sub>** | 0.038 | Productive decomplexation rate, s<sup>-1</sup> |
| **k<sub>stall</sub>** | 0.126 | Rate of stalling (k<sub>cat</sub>/n<sub>app</sub>), s<sup>-1</sup> |
| **n<sub>app</sub>** | 34 | Apparent processivity, Unitless |
| Variables (subscript '0' added to variables indicates initial condition) |  |  |
| **S** | Substrate, g/L |  |
| **S<sub>P</sub>** | Productive binding site, μmole/g |  |
| **S<sub>NP</sub>** | Non-productive binding site, μmole/g |  |
| **S<sub>PT</sub>** | Productive binding capacity of the substrate (maximum number of available productive binding sites per mass of cellulose), μmole/g |  |
| **S<sub>NPT</sub>** | Non-productive binding site capacity of the substrate, μmole/g |  |
| **E** | Free enzyme, μM |  |
| **E<sub>BT</sub>** | Total bound enzyme, μmole/g |  |
| **E<sub>BP</sub>** | Productively bound enzyme, μmole/g |  |
| **E<sub>BNP</sub>** | Non-productively bound enzyme, μmole/g |  |
| **P** | Soluble product, μmole/g |  |
| **CB** | Cellobiose, μM |  |
| **X** | Conversion, % |  |
| **S<sub>P1</sub>** | Productive binding site Population 1, μmole/g |  |
| **S<sub>P2</sub>** | Productive binding site Population 2, μmole/g |  |
| **τ** | Productive binding site lifetime (time basis), hr<sup>-1</sup> |  |
| **γ** | Productive binding site lifetime (conversion basis), %<sup>-1</sup> |  |
| **σ<sub>blocked</sub>** | Fraction of blocked binding sites |  |
| **t** | Time, seconds or hours |  |

**Table 3:** Amplitude and characteristic decay times from double exponential decay fits (Equation 4) to the productive binding capacity of salt washed cellulose (SWC) over time.

|  | Population 1 |  |  | Population 2 |  |  |  |
| --- | --- | --- | --- | --- | --- | --- | --- |
| Cellulose | $S_{P1}$<br>($\mu\text{mole/g}$<br>cellulose)<sup>a</sup> | $\tau_1$ (hr)<sup>a</sup> | Conversion<br>at $\tau_1$ (%) | $S_{P2}$<br>($\mu\text{mole/g}$<br>cellulose)<sup>a</sup> | $\tau_2$ (hr)<sup>a</sup> | Conversion<br>at $\tau_2$ (%) | Reduced<br>$\chi^2$ |
| SC | $4.4 \pm 0.6$ | $0.08 \pm 0.01$ | 40 | $1.3 \pm 0.4$ | $1.4 \pm 0.5$ | 90 | 0.6 |
| BC | $3.7 \pm 0.3$ | $0.05 \pm 0.02$ | 20 | $1.2 \pm 0.6$ | $2.3 \pm 0.1$ | 95 | 0.9 |
| FP | $3.0 \pm 0.5$ | $0.5 \pm 0.1$ | 8 | $0.6 \pm 0.08$ | $30 \pm 5$ | 60 | 0.5 |
| MCC | $2.1 \pm 0.07$ | $0.3 \pm 0.09$ | 2 | $0.3 \pm 0.05$ | $95 \pm 17$ | 50 | 0.002 |
| AC | $1.8 \pm 0.05$ | $0.3 \pm 0.06$ | 4 | $0.2 \pm 0.04$ | $42 \pm 12$ | 40 | 0.001 |

<sup>a</sup> Values represent the best fit parameter obtained from minimization of $\chi^2$ with a 95% confidence interval.

The rate equations for all species in the mechanistic model^11^, including the additional relationship for the rate of change of productive binding capacity (Equation 3) were solved numerically using the ode45 function in MATLAB and with initial conditions matching experimental hydrolysis conditions used in generating the partially hydrolyzed substrates ([S_0_]=0.5 g/L, [E_0_]=2.5 µM, S_P0_ from Figure 4). Hydrolysis was modeled for 120 hours or until 100% conversion was reached. Rate constants derived from the literature are summarized in Table 2, and the justification of these parameters and model development are discussed in greater detail elsewhere^11^. The apparent processivity was held constant for all substrates simulated, and was modeled as unchanging during hydrolysis. It should be noted that the published values of n_app_ vary widely depending on the substrate used^11^, and that conversion-dependent reduction of processivity has been observed for other processive enzyme systems,^62^ and as such, our use of a single unchanging value for n_app_ is a simplification not addressed in this work.

## Results and Discussion

### Initial productive Cel7A binding capacity of cellulose determines the magnitude of the initial burst phase

The productive cellulase binding capacity, [S_PT_], is defined as the number of productive cellulase binding sites per mass of cellulose^31^. The productive Cel7A binding capacity of substrates is determined kinetically from maximum cellobiose release rates (Equation 1) under the premise that product formation is a direct indication of productively bound enzymes. As such, the productive Cel7A binding sites are neither assumed to be cellulose reducing ends per se, nor does this method discount the possibility of ‘endo-initiation’^26^. Rather, the productive Cel7A binding capacity simply counts the maximum number of sites on cellulose where Cel7A has been able to complex and hydrolyze cellulose (Step 1, Figure 2).

As previously observed, despite their similar compositions^31^, the concentration of productive binding sites differed markedly between phosphoric acid swollen filter paper (SC), bacterial cellulose (BC), algal cellulose (AC), filter paper (FP), and microcrystalline cellulose (MCC), where the magnitude of [S_PT_] ranked in the order SC>BC>FP>MCC>AC (Figure 4).

Two of the more recalcitrant celluloses, FP and MCC, were dried during processing, which is thought to cause irreversible aggregation due to hornification and thereby limit cellulase accessibility ^63, 64^. When never-dried SC, BC and AC were air-dried, the productive Cel7A binding capacities dropped dramatically (Figure 4). Extensive acid hydrolysis of cellulose yields highly recalcitrant nanocellulose; here we demonstrate that acid treatment of AC resulted in ∼70% decrease in the productive Cel7A binding capacity.

The order, from high to low productive Cel7A binding capacities for the five cellulosic substrates (SC>BC>FP>MCC>AC), correlates with the conversion reached at the end of the initial burst phase of the hydrolysis curves (Figure 5). Similarly, our previous modeling results indicated that the initial productive cellulase binding capacity limits the extent of initial burst phase kinetics^11, 31^. The productive Cel7A binding capacities in Figure 3, however, are an initial condition and do not indicate the availability of the productive binding sites immediately following the initiation of the reaction. When the productive Cel7A binding capacity is assumed to remain constant throughout the hydrolysis reaction, modeling results over estimate reaction extents^31^. We hypothesized that rate retardations during hydrolysis could be a consequence of the depletion of productive cellulase binding sites on cellulose.

### Productive Cel7A binding sites deplete during the course of cellulose hydrolysis

We determined the evolution of accessible productive Cel7A binding sites over the course of hydrolysis by measuring the productive binding capacities of residual cellulose sampled at various times during the reactions. The partially hydrolyzed cellulose samples were extensively washed with a salt solution to remove surface enzymes and generate ‘salt-washed cellulose’ (SWC). The productive binding capacities of all the celluloses decreased throughout conversion (Figure 5C). The two easily hydrolyzed celluloses with higher initial productive binding capacity, SC and BC, retained higher concentrations of available productive binding sites for the duration of hydrolysis compared to the more recalcitrant FP, MCC and AC.

The productive binding capacity decay trends observed here are in accordance with reported decreases in hydrolysis rates from “restart” experiments with Cel7A^65^ and the decline in specific activity of adsorbed cellulases regardless of cellulose crystallinity^66^. The concentration of cellulase “attack sites” has been estimated from substrate saturating conditions on BC after 30 minutes as 0.96 µmole/g cellulose^22^, and Avicel after 1 hour as 0.22 µmole/g^40^, in good agreement with the values of 1.8 µmole/g for BC at 30 min and 0.4 µmole/g for Avicel at 1 hour reported here. Cellulose is frequently considered an inert material during hydrolysis, yet Figure 5 provides direct evidence that cellulose (as probed by Cel7A) evolves significantly during hydrolysis. Often, a low degree of conversion is used to justify the treatment of cellulose as unchanging in the time frame studied, however, the rapid decay in productive binding capacity at low conversions observed for all celluloses regardless of crystallinity, degree of polymerization, cellulose source or processing history emphasizes the need for careful consideration in selection of cellulosic material and sampling time points in future studies.

A double exponential fit to the productive binding capacities over time suggests two populations of productive binding sites on all celluloses (Table 3). The magnitudes of the productive binding site concentration and lifetime of a given population (S_P_ and τ, respectively) provide insights into the relationship between the productive binding capacity and hydrolysis trends in Figure 5. The first population of productive binding sites, S_P1_, has a higher concentration than S_P2_ for all celluloses —meaning that at lower conversions, more sites from S_P1_ are accessible to Cel7A than S_P2_. The fact that τ_1_ is significantly shorter than τ_2_, indicates that a majority of the sites in Population 1 are depleted before significant enzyme action has occurred on Population 2. The lifetime of the larger population of productive binding sites, τ_1_, extends to a much greater conversion (∼20-40%) for SC and BC than for the more recalcitrant celluloses FP, MCC and AC, where these sites are, on average, depleted before 10% conversion. This, along with the ranking of the magnitudes of S_P1_ for each cellulose, is reflected in the extent and duration of burst phase kinetics for the celluloses, which rank in order of SC>BC>FP>MCC>AC.

Population 2—as the smaller, more slowly decaying population of productive binding sites—governs longer-term hydrolysis trends once a majority of the productive binding sites from the more accessible Population 1 are consumed. The initial concentration of this second population of sites is ∼30% of the total number of productive binding sites for rapidly hydrolyzed substrates, while it is about 15% of total binding sites for the less readily hydrolyzed celluloses. The magnitude of S_P2_ correlates with hydrolysis rates at higher conversions, as well as overall conversions, where SC>BC>FP>MCC>AC. As productive binding sites from Population 1 are depleted, it would make sense that those remaining in Population 2 become increasingly rate limiting, hence the correlation of S_P2_ with late term hydrolysis rates. Furthermore, X at τ_2_ correlates with overall conversion, which is reasonable as no further hydrolysis can occur after [S_PT_] decays to zero.

Cellulose hydrolysis has been described by two parallel first order reactions of “easy” and “difficult” to hydrolyze cellulose in the literature.^50, 67, 68^. While the fits to the time-evolution of the productive Cel7A binding capacities are empirical and true physical meaning cannot be ascribed to S_P1_ and S_P2_, we conjecture that Population 1 represents initially accessible surface binding sites and is a substrate related feature contributing to the slowdown of the initial burst in hydrolysis. The second population of productive binding sites we interpret as surface sites that become exposed when those around them are removed, and the uncovering of these sites occurs at a more slowly decaying rate during the “steady state” phase of hydrolysis. In the first 30 minutes of hydrolysis, a large magnitude, short lived population, and a smaller magnitude, longer lifetime population of “hydrolysis initiations” measured under single turnover conditions where each bound cellulase is allowed to perform only a single processive run has previously been reported^24^. As each processive run must start at a productive binding site, these populations are essentially another estimate for the evolving productive binding capacity, and agree with the data shown here.

The sustained lifetime of Population 1 and the relatively high magnitude of S_P2_ for BC are in agreement with the hypothesis that the tendency of this cellulose to fibrillate upon treatment with Cel7A exposes new productive binding sites, as is the rapid depletion of available sites on AC, which remains tightly packed after enzyme treatment^56^. However, lacking in a “chain exposure” model is a physical explanation of what makes the sites in Population 1 more accessible for substrates like BC and SC when compared to FP, MCC and AC. If the only limitation on accessibility is exposure as a surface site, as chain ablation and surface area models assume^8^, then we would expect a dependency of productive binding site concentrations on surface area. However, the specific surface areas reported for MCC as 0.96-2.4 m^2^/g^69–71^, AC as 64.3-94.7 m^2^/g ^69, 70^, and BC as 39.2 m^2^/g^71^, do not correlate with the initial productive binding capacity of MCC and AC, nor the larger productive binding capacity of BC.

### ‘Irreversibly bound’ surface enzymes contribute to the depletion of productive binding sites

Although Figure 5 suggests that hydrolysis by Cel7A causes the decline in available productive binding sites on the cellulosic substrates, an alternate or confounding possibility is that the productive binding sites are simply becoming obstructed by tightly or irreversibly bound Cel7A. This question was addressed by generating a parallel set of partially hydrolyzed cellulose samples that were additionally treated with a protease to remove any residual Cel7A on the substrate surface. Analysis of salt-washed cellulose (SWC) and protease-treated cellulose (PTC) samples by sFTIR showed that a small portion of residual enzymes remained on the surface of SWC, while the additional protease treatment completely removed any surface protein (Figure S2 in SI). The productive binding capacities of the PTC showed similar decays over the time course as observed with SWC (Figure 5), but retained higher overall magnitudes of productive binding sites throughout hydrolysis (fits are provided in Table S1 in SI). This implies that while decay trends are substrate related in origin, irreversibly adsorbed enzymes further contribute to rate limitations, as has been suggested before^16, 20, 21, 32, 33, 72, 73^.

To isolate the substrate contribution to rate slowdown, we compared the estimated number of accessible productive binding sites for enzyme-free PTC and residual enzyme containing SWC (Figure 6). We observed a systematic increase in productive binding capacity upon full removal of surface associated enzymes. The fraction of productive binding sites blocked by bound enzyme, σ_Blocked_ was calculated as:

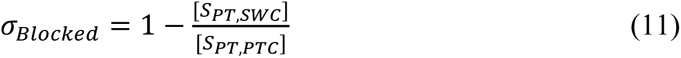

and was statistically invariant as conversion increased. The time-averaged fraction of blocked sites is summarized in Table 4.

**Figure 6:**
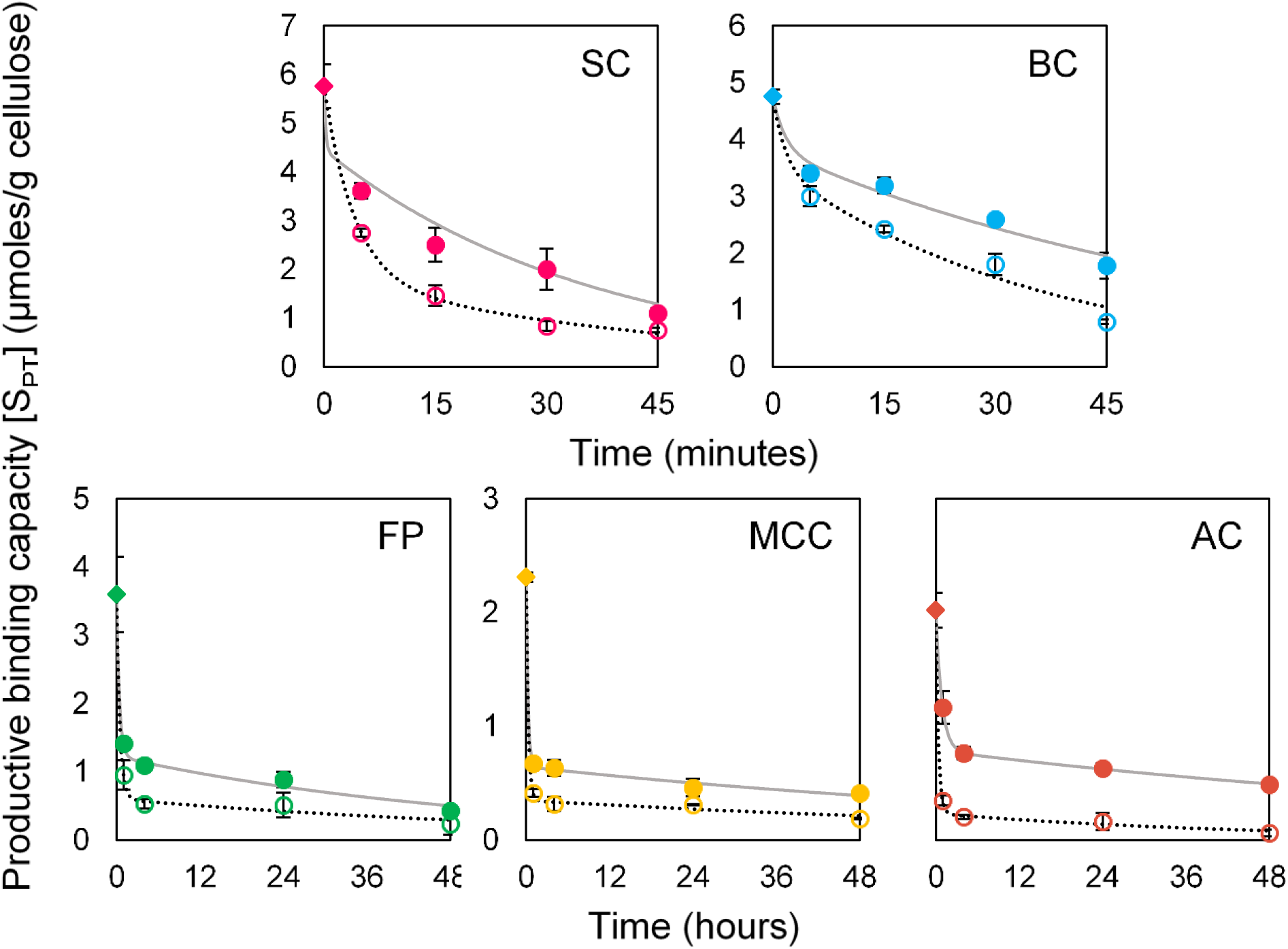
Comparison of the productive Cel7A binding capacity of salt washed cellulose (SWC, open markers) and protease treated cellulose (PTC, filled markers) as a function of time. Lines represent fits by Equation 4 for SWC (dotted lines), and PTC (solid line). Fitting parameters are summarized in Table 3 and S1 in SI.

**Table 4:**
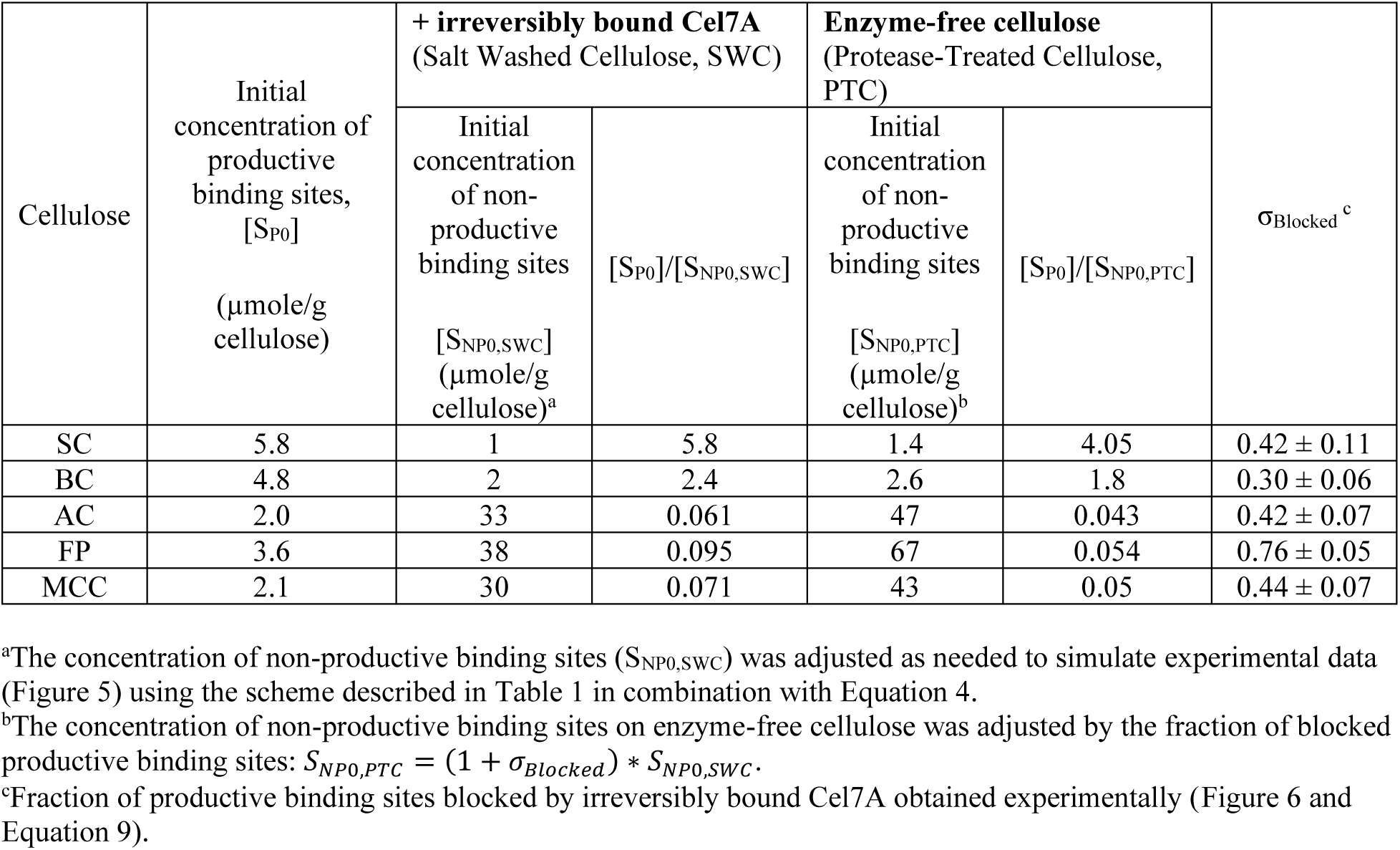
Summary of the initial concentration of productive binding sites ([S_P0_]), non-productive binding sites ([S_NP0_]), and initial ratio of sites ([S_P0_]/[S_NP0_]). The subscript SWC or PTC indicates salt washed cellulose with irreversibly bound enzyme or enzyme-free protease-treated cellulose, respectively. The fraction of sites blocked by irreversibly bound enzyme on salt washed cellulose calculated with Equation 9 is also tabulated (σ_Blocked_).

While the reversibility of cellulase binding to cellulose and its role in hydrolysis rate retardation remains contradictory^21, 22, 72–80^, our data strongly suggest that some Cel7A resist removal by extensive salt washing, and remain tightly bound to and block significant fractions of the productive binding sites on celluloses (Table 4). We confirm that this residual protein is catalytically inactive from flat biosensor readouts indicating no cellobiose production. The onset of enzyme blockage of accessible productive binding sites occurs rapidly (site blockage has already reached a maximum at the first time point measured), but the percentage of blocked sites does not change throughout hydrolysis. The unchanging obstruction of sites suggests that surface enzymes may contribute to retardation of the burst phase to steady state rates, but does not explain further rate limitations occurring at greater conversions. This concept has been captured by enzyme centric models which incorporate non-productive binding, but treat the substrate as inert.^8^ Inspection of productive binding capacity trends for PTC shows that even without the presence of residual cellulases, the concentration of productive binding sites available for Cel7A still depletes by a double exponential decay (Figure 6 and Table S1 in SI), indicating that substrate related evolution governs rate limitations at greater conversions.

### Depletion of the productive binding sites explains overall hydrolysis kinetics

Incorporating the time dependence of productive binding site concentrations successfully simulates experimental cellulose hydrolysis kinetics by mechanistic modeling. The time evolution of the substrate, i.e. the double exponential depletion of productive binding sites as a function of time (Equation 3, Table 3) was incorporated with the kinetic mechanism (Equations 4-8, Table 2) to simulate Cel7A hydrolysis of SC, BC, FP, MCC and AC. The simulations successfully captured hydrolysis trends throughout the extended time course (Figure 5); namely, the curvature of experimental data both during the burst phase and at longer hydrolysis phases are accurately simulated. For example, MCC and AC exhibit similar conversion trends during the first 24 hours of hydrolysis, but AC plateaus more sharply to reach a lower overall degree of conversion, which is accurately simulated by our model. These trends are reflected by the similar decrease in productive binding capacity to 24 hours (∼10% conversion of both substrates), with a lower long-term binding capacity resulting for AC (Figure 5).

The initial concentrations of non-productive binding sites (S_NP0,SWC_, Table 4) were adjusted for each cellulose to accurately simulate the experimental hydrolysis time courses in Figure 5. The easily digestible celluloses, SC and BC had lower concentrations of non-productive binding sites while the recalcitrant celluloses (FP, MCC and AC) had an order of magnitude higher concentrations of non-productive binding sites. The ratio of initial productive to non-productive binding capacities, S_P0_/S_NP0,SWC_, tracked with the overall digestibility of the substrates (SC>BC>FP>MCC>AC), with values around 2 – 6 for the more hydrolysable substrates and <0.1 for the more recalcitrant celluloses. Higher initial ratios of productive to non-productive binding sites could be a quantitative indicator of higher cellulase accessibility to a given cellulosic substrate.

A 10-fold decrease in enzyme loading predictably resulted in decreased hydrolysis rate and conversion of AC (Figure 7A). Most interestingly, the hydrolysis time course of AC by the decreased enzyme loading of 0.25 µM could be successfully simulated by only increasing τ_1_ from 0.35 hr to 1.4 hr and keeping all other parameters the same (Figure 7A). This phenomenon can be explained by a productive binding site exposure model, where we would expect that the total enzyme concentration have a large effect on how quickly productive binding sites are depleted when they are in relative abundance (or a more accessible), i.e., the productive binding sites in Population 1. In this line of reasoning, as the enzyme loading is decreased, the lifetime of the initially depleted fraction of productive binding sites, τ_1,_ should get longer. In the substrate limited regime (higher conversions/times), the productive binding sites of Population 2 will only be depleted as quickly as they are exposed. The rate of exposure of these sites would depend largely on inherent substrate properties, and enzyme loading should have little effect on τ_2_. Our model predicts lower concentrations of productively bound enzyme (Figure 7B) with lower total enzyme loading, indicating lower occupancy of productive binding sites at lower total enzyme loading.

**Figure 7:**
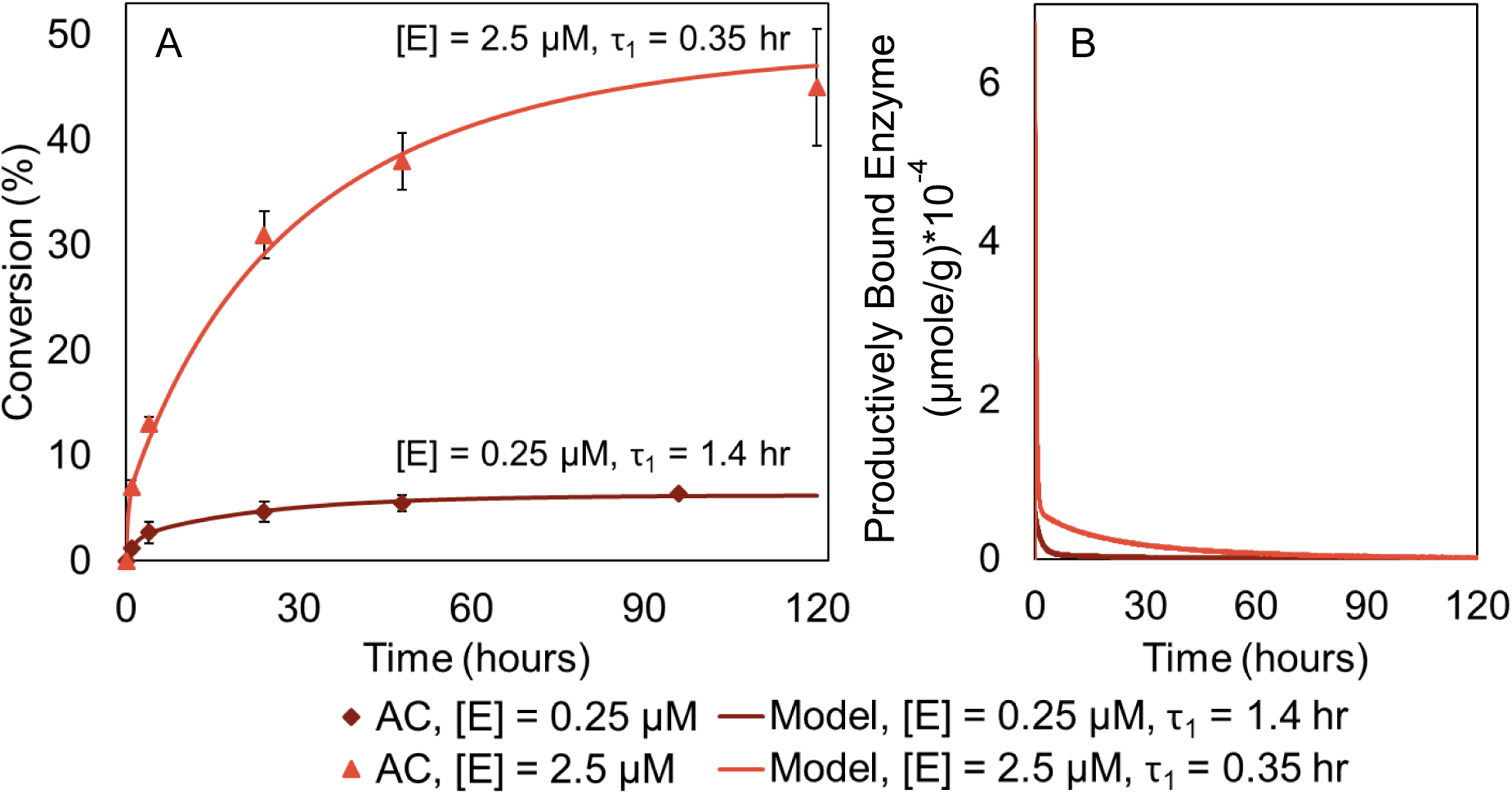
Simulation of hydrolysis of AC with TrCel7A at various enzyme loadings. A) The model predicted conversion (solid lines) and experimentally determined conversion (solid markers) of AC at 0.25 µM (dark red diamonds) and 2.5 µM (red triangles) enzyme loading. B) The simulated concentration of productively bound enzyme of TrCel7A on AC at 0.25 µM (dark red line) and 2.5 µM (red line) enzyme loadings.

The removal of irreversibly bound Cel7A exposed additional non-productive binding sites on all the celluloses. Increasing the initial concentrations of non-productive binding sites (S_NP0,PTC_) by the percentage of sites blocked by irreversibly bound enzymes (Table 4) when the simulation was conducted using the productive binding capacity decay of enzyme-free cellulose (d[S_P,PTC_]/dt, Table S1) successfully simulated experimental data of BC hydrolysis (‘Adjusted PTC’ curve in Figure 8). Conversely, when the initial concentration of non-productive binding sites was not adjusted to compensate for blocked sites, the simulation over-estimated cellulose conversion by extending the burst phase of the reaction (‘Unadjusted PTC’ curve in Figure 8). These simulations suggest that irreversible enzyme binding that block potential productive binding sites reduces overall conversion potential of the cellulose by truncating or limiting the burst phase of the hydrolysis reaction.

**Figure 8:**
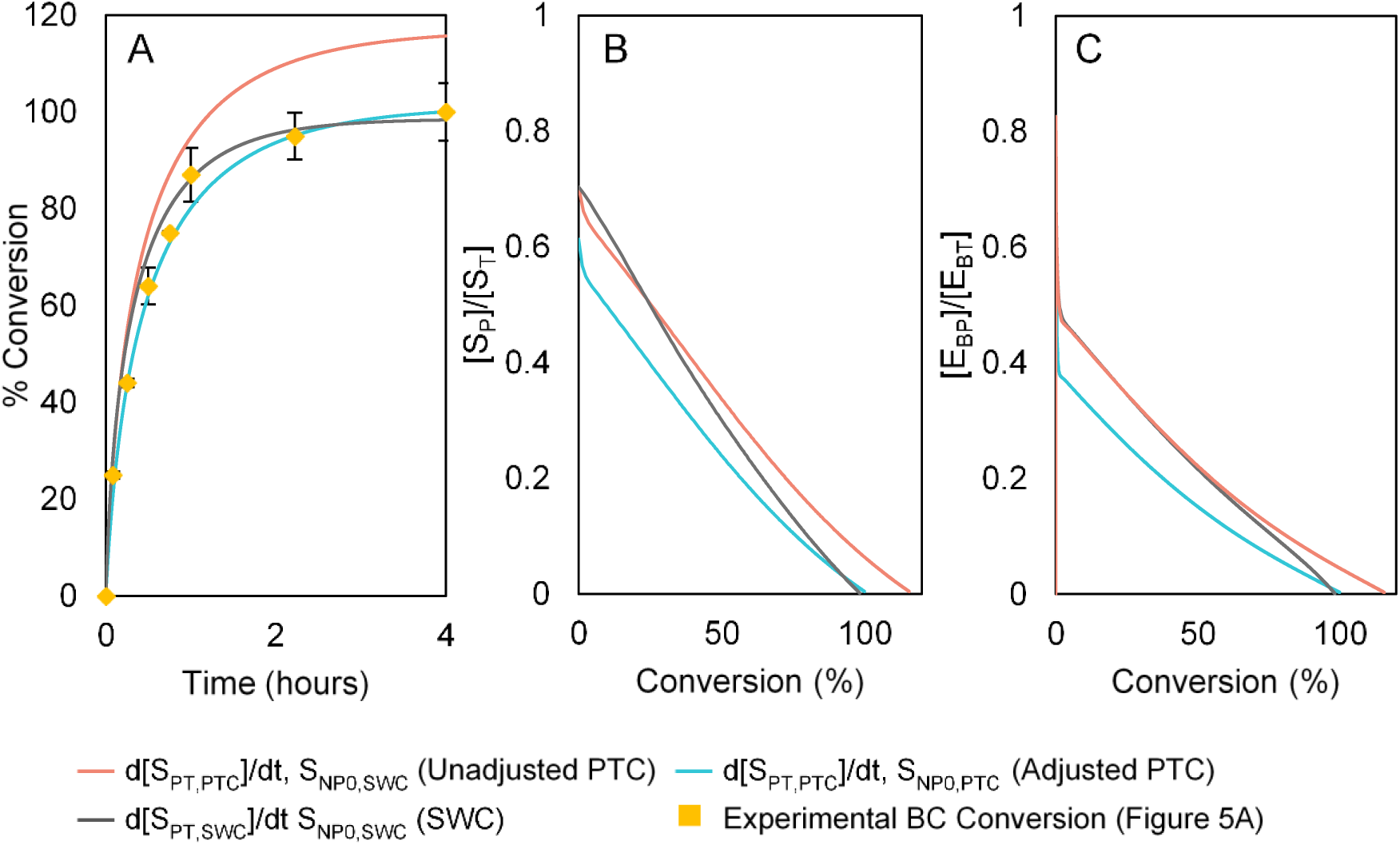
A) The hydrolysis time course of BC simulated using initial concentration of non-productive binding sites, S_NP0,SWC_ and S_NP0,PTC_. B) Simulations of the fraction of productive binding sites on cellulose, and C) fraction of productively bound enzyme plotted versus conversion of BC.

### Decreasing non-productive binding sites promotes productive binding leading to increased hydrolysis rates

A closer look at the binding sites and bound enzyme distributions (Figure 8B and C) reveals that a higher fraction of productive binding sites ([S_P_]/[S_T_], Figure 8B) results in a higher fraction of productively bound enzyme ([E_BP_]/[E_BT_], Figure 8C), leading to faster hydrolysis rates both in the initial burst phase and at longer reaction times (Figure 8A). Initially, the presence of irreversibly bound enzymes blocking productive binding sites has little influence on either the overall fraction of productive binding sites or the fraction of productively bound enzymes. This can be seen by similar ratios of productive binding sites and productively bound enzymes between SWC with irreversibly bound enzyme (‘SWC’ in Figure 8) and enzyme-free cellulose (‘Unadjusted PTC’ in Figure 8). A slower depletion of productive binding sites on enzyme-free cellulose is responsible for the longer burst and higher overall conversion reached in Figure 8A. Thus, we see that preventing irreversible enzyme binding that results in the loss of productive binding sites maintains a higher fraction of productively bound enzyme and helps to overcome cellulose recalcitrance.

Bacterial cellulose, one of the easily hydrolysable celluloses, had a relatively high initial ratio of productive to non-productive binding sites compared to the recalcitrant celluloses; e.g. [S_P0_]/[S_NP0_] = 2.4 and 0.06 for BC and AC, respectively (Table 4). Simulating a scenario where the [S_P0_]/[S_NP0_] ratio of BC was reduced to 0.06 (by increasing the initial concentration of non-productive binding sites, S_NP0_ from 2 to 80 µmoles/g) greatly limited hydrolysis rates of BC, where the maximum achievable conversion dropped to ∼10 % from nearly 100% (Figure 9A). Likewise, when the [S_P0_]/[S_NP0_] ratio for AC was increased 40-fold to 2.4 by decreasing [S_NP0_] from 33 to 0.85 µmole/g, the maximum conversion increased from ∼50% to 120% (Figure 9B). The conversion increase was primarily due to an increase in burst phase hydrolysis rates due to an overall increase in productive binding by the enzymes (Figure S7) even though the concentration of productive binding sites was also not changed in this simulation. Increasing concentrations of non-productive binding sites alone (i.e. with no change in the concentration of productive binding sites) appear to both increase non-productively bound enzymes and decrease productively bound enzymes. In fact, Figure 9A and B demonstrate that the hydrolyzability of cellulose changes dramatically simply by changing the relative concentration of non-productive binding sites.

**Figure 9:**
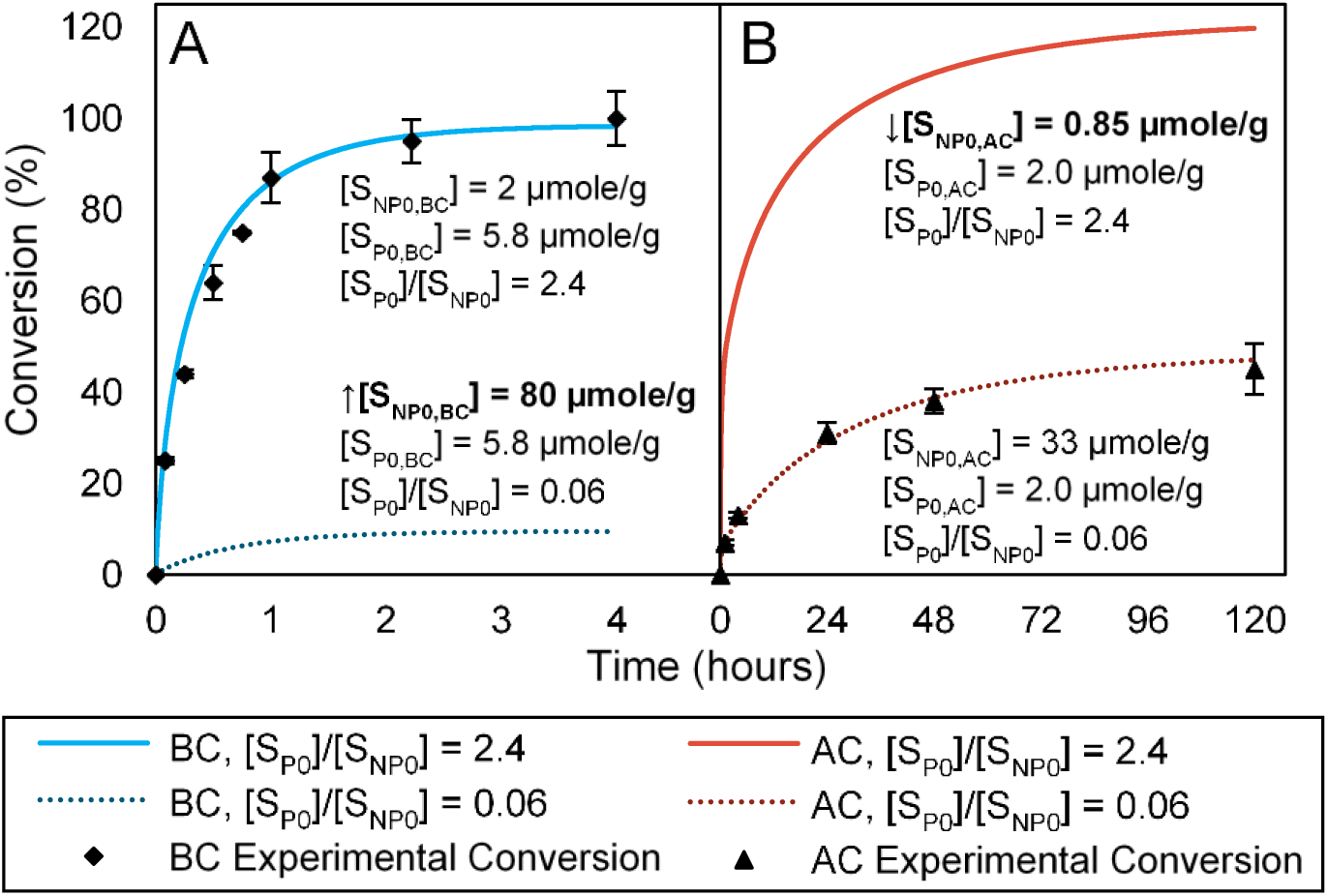
A) Simulated conversion of BC with [S_P0_]/[S_NP0_] ratio of 2.4 and 0.06 over time (alongside experimental conversion of BC from Figure 5A); B)simulated conversion of AC with [S_P0_]/[S_NP0_] ratio of 2.4 and 0.06 with respect to time (alongside experimental conversion of AC from Figure 5A); model predictions for the concentration of non-productively bound enzyme with respect to conversion of BC (C) and AC (D) with [S_P0_]/[S_NP0_] ratios of 2.4 and 0.06; model predictions of the concentration of productively bound enzyme with respect to conversion of BC (E) and BC (F) with [S_P0_]/[S_NP0_] ratios of 2.4 and 0.06; inset in (F) is a zoomed plot of the concentration of non-productively bound enzyme on AC with [S_P0_]/[S_NP0_]=0.06.

## Summary and Conclusions

The discrepancy between reported total binding capacities of cellulose^20,22,40,81^and the productive binding capacities measured here highlights that total surface saturation and productive site saturation are not necessarily interchangeable. In addition, we have shown that the productive binding capacity of cellulose evolves throughout hydrolysis, and that in some instances, treatment of the substrate as inert in heterogeneous interfacial catalysis may inaccurately capture reaction mechanisms. In the context of selecting appropriate reaction conditions to maximize hydrolytic efficiency, estimates for [S_PT_] throughout hydrolysis can aid in tailoring substrate processing to maximize productive binding site availability initially and during hydrolysis, as well as informing process design to minimize the contribution of inactive surface enzymes in rate limitation.^13, 20, 35, 50, 81, 82^

From this study, we have found that the extent to which the initial rate can be maximized and sustained determines overall rates and extents of conversion of cellulose. The results shown here suggest that changes in accessibility of the substrate are the primary source of hydrolysis slowdown. We found that increasing the ratio of productive- to non-productive binding sites promotes hydrolysis, while maintaining an elevated productive binding capacity throughout conversion is key to preventing hydrolysis slowdown. The methods developed here serve as an excellent tool to analyze the effectiveness of substrate pretreatment and processing in the context of increasing cellulose accessibility. Measurements of the productive binding capacity can also be used to evaluate the success of protein engineering of both monocomponent and synergistic enzyme systems, and a more detailed understanding of mechanisms of both productive and non-productive binding can inform more targeted protein modifications. Although further study is needed to identify the physical meaning of productive and non-productive binding sites, we conclude that the initial and evolving concentrations of binding site populations on the surface of an insoluble substrate such as cellulose plays a significant role in limiting reaction mechanisms of heterogeneous enzyme catalysis.

## Supporting Information Available

Information further detailing experimental methods and results, fitting parameters, and model simulations is included in Supporting Information.

## Acknowledgements

We would like to acknowledge Novozymes, Inc. for providing cellobiose dehydrogenase and Alex Hitomi for assistance in preparation of untreated and acid hydrolyzed algal cellulose. This research used resources of the Advanced Light Source, which is a U.S. Department of Energy (DOE) Office of Science (OS) User Facility under contract no. DE-AC02-05CH11231. This work was supported in part by the DOE BER Bioimaging Science Program (Award No. DE-SC0019228), the ALS Doctoral Fellowship in Residence Program and the DOE OS Office of Workforce Development for Teachers and Scientists, Office of Science Graduate Student Research (SCGSR) program. The SCGSR program is administered by the Oak Ridge Institute for Science and Education for the DOE under contract number DE-SC0014664. This work made use of the BSISB Resource supported by DOE/BER under Contract No. DE-AC02-05CH11231.

## TOC Graphic

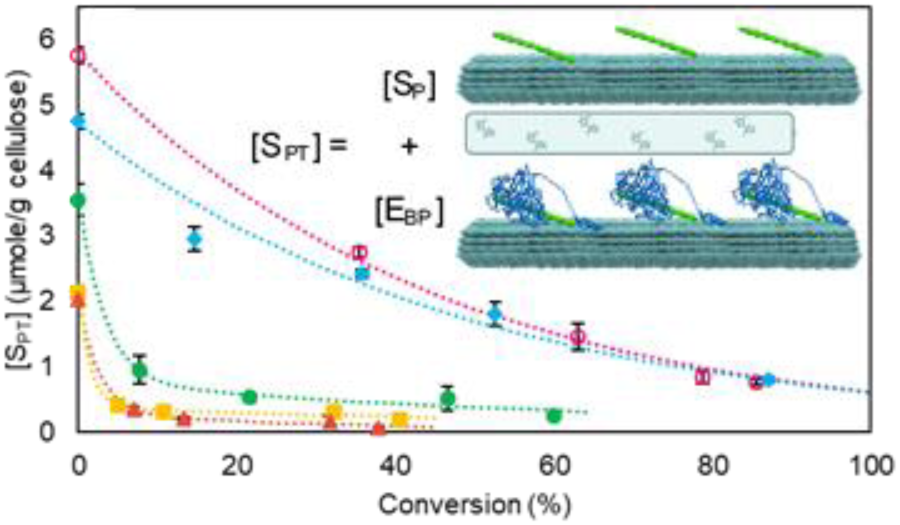

