## Supporting Information for "The Role of Evolving Interfacial Substrate Properties on Heterogeneous Cellulose Hydrolysis Kinetics"

### Solids loss in centrifuge tubes

Unhydrolyzed cellulose was washed following the procedure for cellulose washing in the main text using MilliQ water in place of 0.5 M NaCl. The concentration of unwashed and washed cellulose was measured by Anthrone assay using glucose standards in triplicate as previously described[1, 2]. The solids loss was determined to be negligible as shown in Figure S1.

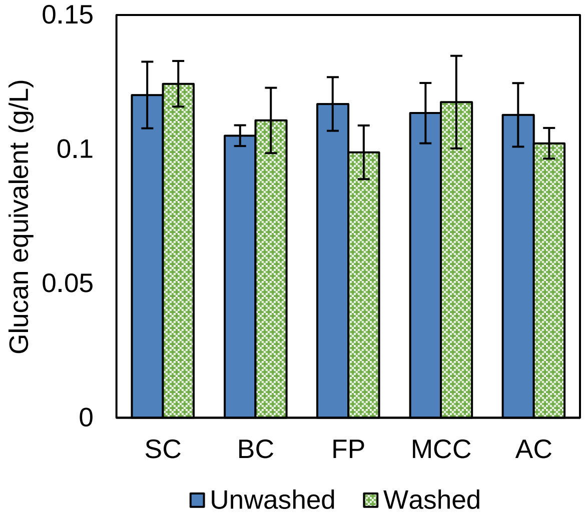


*Figure S1: Cellulose concentration determined by Anthrone assay before and after washing in centrifugation filter tubes of swollen cellulose (SC), bacterial cellulose (BC), filter paper (FP), microcrystalline cellulose (MCC) and algal cellulose (AC).*

### <u>Synchrotron Fourier Transform Infrared Spectroscopy (sFTIR) of Cellulose</u>

A small aliquot (~1-2 uL) of each sample was dried on a ZnSe lens and cellulose fibrils were located using a 32X objective with an optical microscope. Infrared transmission spectra were acquired with a synchrotron-based FTIR microscope on Beamline 1.4.3 at the Advanced Light Source (ALS) in the 650–4000 $cm^{-1}$ range with 4 $cm^{-1}$ resolution using a KBr beamsplitter and a mercury cadmium telluride (MCT-A) detector. Spectra were normalized to a background obtained on a blank area and vector normalized for comparison.

Unwashed partially hydrolyzed cellulose was produced by terminating the reaction by filtration and resuspending in MilliQ water at 4 $^oC$. sFTIR spectra of unhydrolyzed, unwashed partially hydrolyzed, NaCl washed and Protease treated algal cellulose are shown in Figure S2 and are representative of enzyme binding trends observed for all celluloses studied. Peaks at ~1500-1700 corresponding to amide peaks indicating the presence of protein are apparent on unwashed and NaCl washed celluloses, which are absent on unhydrolyzed and protease treated cellulose.

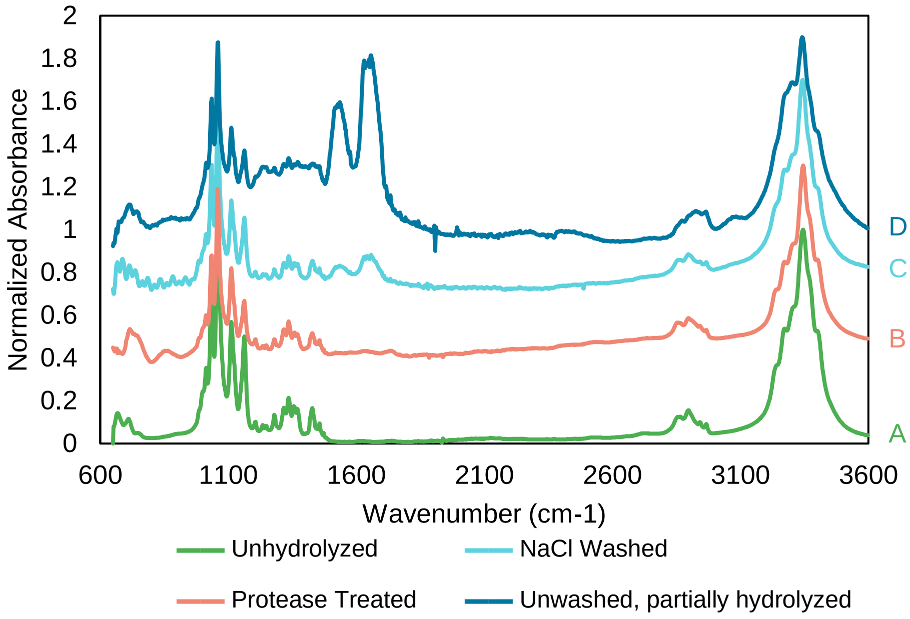


*Figure S2: sFTIR spectra of A) Unhydrolyzed; B) Protease Treated; C) NaCl washed; and D) Unwashed, partially hydrolyzed algal cellulose which is representative of spectra observed for all celluloses studied.*

### Biosensor calibration

A cellobiose dehydrogenase (CDH) based biosensor was calibrated immediately preceding and following each measurement by stepwise addition to a final cellobiose concentration of 40 µM. Probes typically had a linear response up to ~30 µM, and a representative calibration curve is shown in Figure S3.

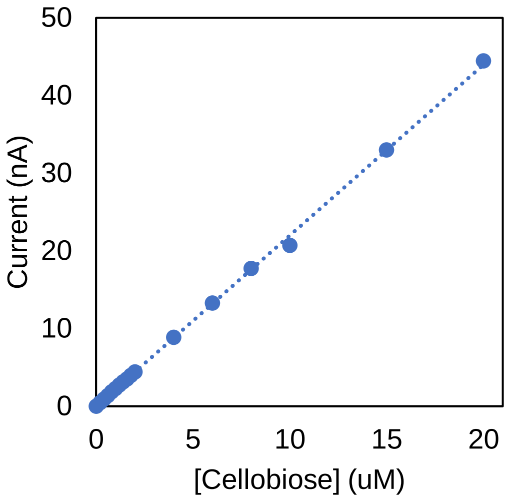


*Figure S3: Calibration curve of CDH biosensor using stepwise additions of cellobiose.*

### Validation of biosensor measurements

Aliquots from reactions conducted during measurement by biosensor were stopped at 30 seconds by filtration and measured with high performance anion exchange chromatography with pulsed amperometric detection (HPAE-PAD) to verify the cellobiose concentrations reported by the biosensor. Comparison of the measured cellobiose release after 30 seconds for reactions of 0.5 g/L of cellulose with 130 μmole/g Cel7A by biosensor and HPAEC-PAD are shown in Figure S4, and are not significantly different.

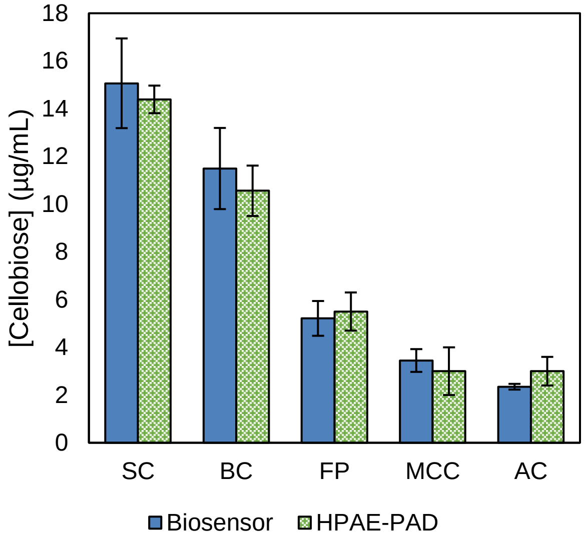


*Figure S4: Comparison of sugar release detected by an amperometric CDH biosensor and HPAE-PAD after 30 seconds of hydrolysis of 0.5 g/L of cellulose with 130 μmole/g Cel7A.*

From our previous work, we expected to measure values of productive binding in the range of 0.1-10 μmoles/g of cellulose[3]. Using $k_{cat}$=5.4 $s^{-1}$ we then expect that our experiments required that the sensor accurately measure a rate of cellobiose release, d[CB]/dt, accurately in the range of 0.02-3 μM/s. To simulate these slopes, we injected cellobiose to final concentrations ranging from 0.1-20 μM, corresponding to maximum hydrolysis rates of ~0.02-6 μM/s, as shown in Figure S5A (0.1 and 0.5 μM additions have been omitted for clarity). The measured rates and the corresponding [$E_{BP}$] estimates were linear for cellobiose additions from 0.1-20 μM, as shown in Figure S5B, confirming that the experimental system accurately captures initial hydrolysis rates in ranges relevant to experiments conducted here.

Real-time progress curves were measured on each cellulose at enzyme/substrate ratios ranging from 5 μmol/g cellulose to 130 μmol/g cellulose. A representative product curve and the estimation of the concentration of productively bound enzyme from the maximum hydrolysis rate is shown in Figure S5C. The total number of available productive binding sites (the productive binding capacity, $S_{PT}$) was estimated at saturation concentrations of productively bound enzyme (occurring when no further increase of hydrolysis rates occurs with higher enzyme loading), as shown in Figure S5D. The saturation trend shown in Figure S5D was observed for all cellulose types studied here.

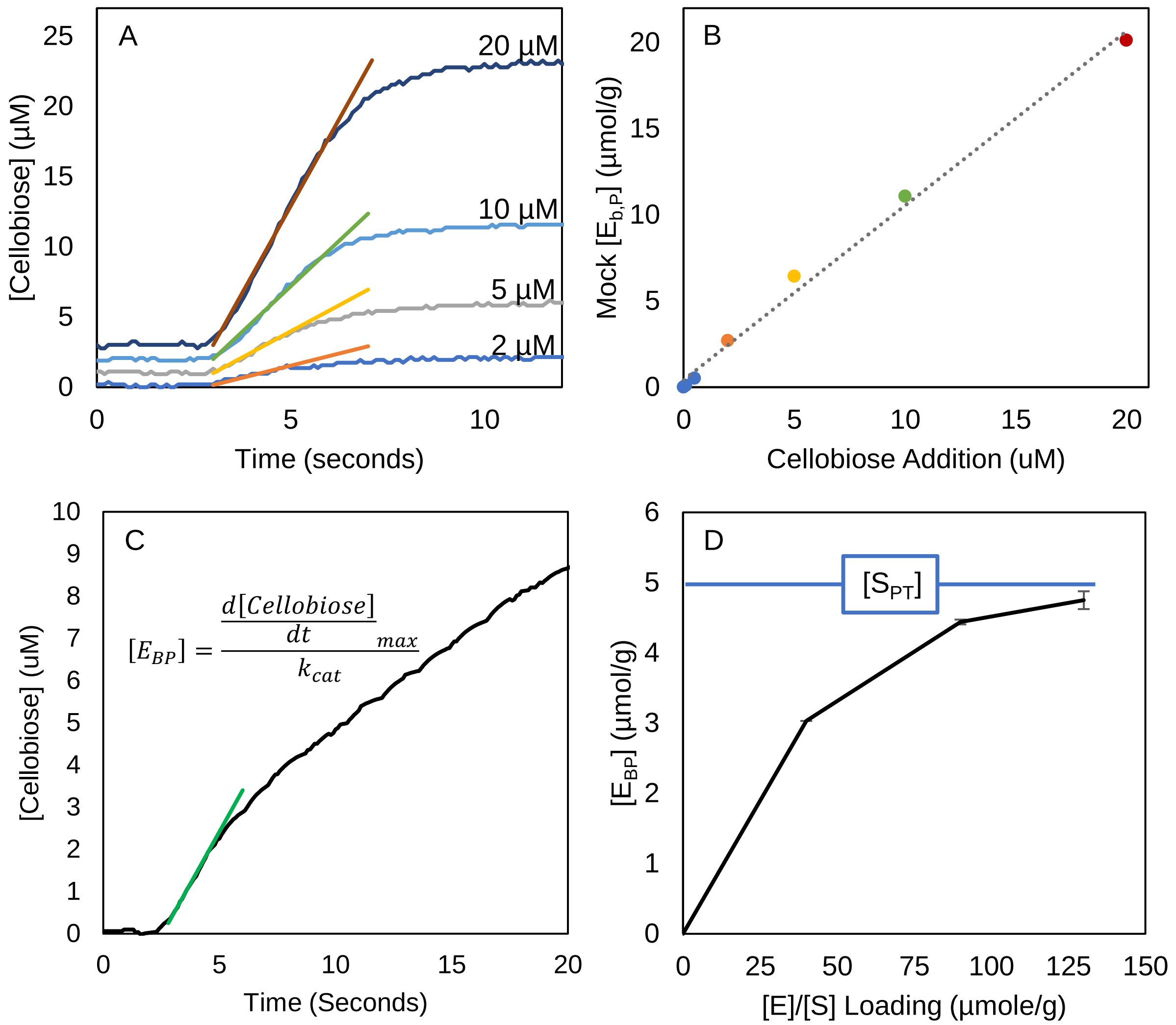


*Figure S5: A) Estimation of "Mock [$E_{BP}$]" from increasing cellobiose additions; B) Plot of estimated Mock [$E_{BP}$] versus cellobiose addition; C) Representative curve of estimation of [$E_{BP}$] from cellulose hydrolysis data; D) Representative curve of the estimation of the productive binding capacity, $S_{PT}$ from saturation curves of productive binding sites.*

The ability of a CDH biosensor to capture initial hydrolysis rates of cellulose by Cel7A has been previously justified[4, 5], and validation of the system's capability to capture reaction rates relevant to the reaction conditions used here (~0.03-6 µM/s) is shown here. The agreement between previously reported initial productive binding capacities of cellulose estimated from fits to 3 timepoints within the first 30 seconds of hydrolysis[3] and from the real time data here validates the application of a CDH biosensor to obtain initial hydrolysis rates for use in Equation 1 (main text).

The characteristic sigmoidal curve with rate maximums occurring around 3-7 seconds after enzyme addition with subsequent rate retardation observed at lower enzyme to substrate loadings ([E]/[$S_0$]) and higher overall solids loadings[5-8] was also observed here at substrate saturating conditions (high [E]/[$S_0$] and low solids), as shown in Figure S5A and C. Cruys-Bagger et. al reported an apparent disappearance of the burst at low substrate concentrations

(0.25 g/L)[4], but we propose that the burst observed here at low solids is possibly due to the much higher enzyme loadings utilized.

Using an estimate of $k_{cat}$~5.4 $s^{-1}$, it follows that the use of a 2 second interval to fit $d[CB]/dt_{max}$ allows for ~10 catalytic events and should be before the low estimate for apparent processivity (13 catalytic events)[7, 9], where enzymes reach the end of the first obstacle free path. Further evidence the fitting region selected was prior to any slowdown from enzyme dissociation or blockage comes from the good linear fit of the data in this time range shown in Figures S5A and C. Product inhibition can be assumed to be negligible in these experiments because the maximum sugar concentrations measured here (~15 uM) are much less than reported inhibition constants for Cel7A on insoluble substrates.[10-15]

### Fitting parameters

Data for the productive binding capacity was fit in the main text using Equation 2 as a function of time (Figure 5, main text) or as a function of conversion (Figure 4, main text) using cftool in MATLAB and Equation S1,

$$[S_{PT}] = S_{P1}\exp\left(-{}^{X}\!/_{\gamma_1}\right) + S_{P2}\exp\left(-{}^{X}\!/_{\gamma_2}\right) \qquad \text{(S1)}$$

where $S_P$ is the concentration of productive binding sites (μmol/g), γ is the decay constant (%) and X is conversion (%). The fitting parameters are tabulated in Table S1 for all cases. Comparison of the productive binding capacities of SWC and PTC as a function of conversion are shown in Figure S6.

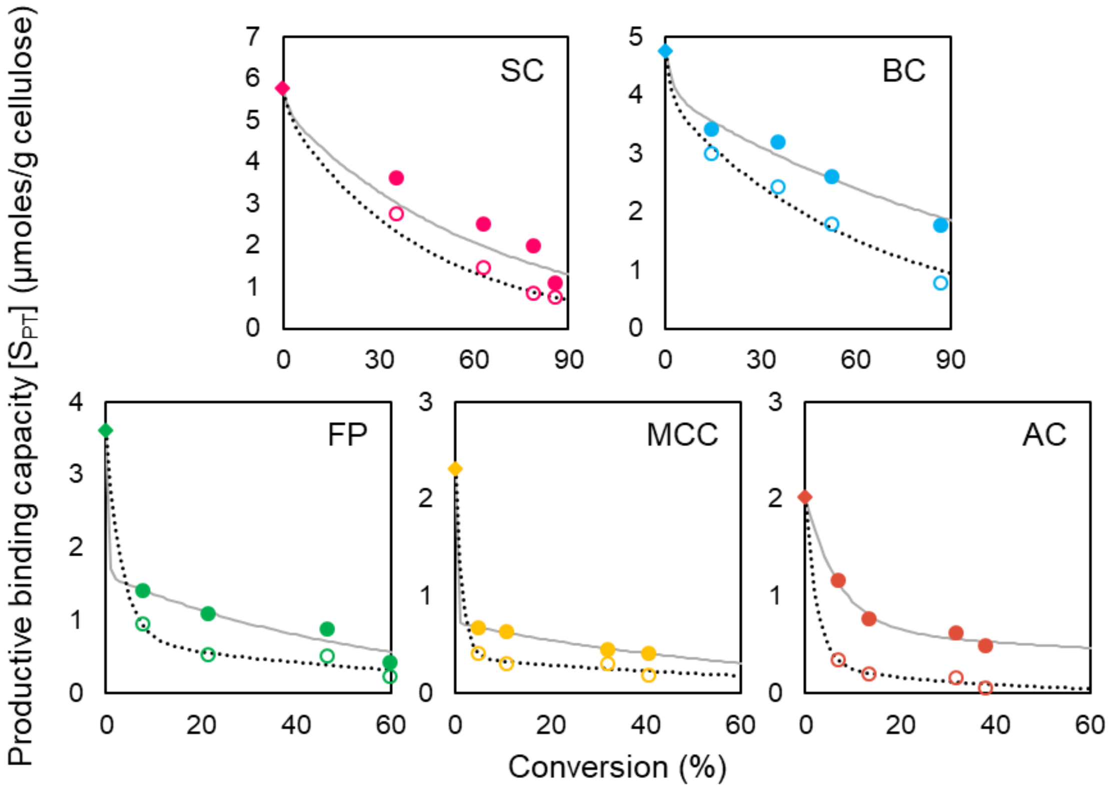


*Figure S6: Comparison of the productive binding capacity of salt washed cellulose (SWC, open markers) and protease treated cellulose (PTC, filled markers) as a function of conversion. Lines represent fits by Equation S1 for SWC (dotted lines), and PTC (solid line). Fitting parameters are summarized in Table S1.*

*Table S1: Fitting Parameters for the productive binding capacity of salt washed cellulose (SWC) and protease-treated cellulose (PTC) as a function of time and conversion. Values represent the best fit parameter from minimization of $\chi 2$ with a 95% confidence interval.*

| SWC as a function of time | | | | | |
|---|---|---|---|---|---|
| Cellulose | SC | BC | FP | MCC | AC |
| $S_{P1}$ (µmole/g cellulose) | 4.4 ± 0.6 | 3.7 ± 0.3 | 3.0 ± 0.5 | 2.1 ± 0.07 | 1.8 ± 0.05 |
| $S_{P2}$ (µmole/g cellulose) | 1.3 ± 0.4 | 1.2 ± 0.6 | 0.6 ± 0.08 | 0.3 ± 0.05 | 0.2 ± 0.04 |
| $\tau_1$ (hr) | 0.08 ± 0.01 | 0.05 ± 0.02 | 0.5 ± 0.1 | 0.3 ± 0.09 | 0.3 ± 0.06 |
| $\tau_2$ (hr) | 1.4 ± 0.5 | 2.3 ± 0.1 | 30 ± 5 | 95 ± 17 | 42 ± 12 |
| Reduced $\chi^2$ | 0.9 | 0.9 | 0.5 | 0.002 | 0.001 |
| PTC as a function of time | | | | | |
| Cellulose | SC | BC | FP | MCC | AC |
| $S_{P1}$ (µmole/g cellulose) | 3.9 ± 0.8 | 3.7 ± 0.3 | 2.4 ± 0.6 | 1.7 ± 0.08 | 1.2 ± 0.2 |
| $S_{P2}$ (µmole/g cellulose) | 1.8 ± 0.9 | 1.1 ± 0.3 | 1.2 ± 0.1 | 0.6 ± 0.07 | 0.8 ± 0.06 |
| $\tau_1$ (hr) | 0.04 ± 0.04 | 0.03 ± 0.25 | 0.4 ± 0.1 | 0.3 ± 0.1 | 0.84 ± 0.3 |
| $\tau_2$ (hr) | 0.7 ± 0.1 | 0.7 ± 0.2 | 47 ± 6 | 111 ± 20 | 100 ± 22 |
| Reduced $\chi^2$ | 0.6 | 3.9 | 2.0 | 0.6 | 0.1 |
| SWC as a function of conversion | | | | | |
| Cellulose | SC | BC | FP | MCC | AC |
| $S_{P1}$ (µmole/g cellulose) | 5.2 ± 0.2 | 3.9 ± 0.5 | 2.9 ± 0.6 | 1.9 ± 0.7 | 1.7 ± 0.5 |
| $S_{P2}$ (µmole/g cellulose) | 0.5 ± 0.08 | 0.8 ± 0.6 | 0.7 ± 0.2 | 0.4 ± 0.1 | 0.3 ± 0.05 |
| $\gamma_1$ (%) | 2.0 ± 0.02 | 2.0 ± 0.5 | 3.3 ± 0.1 | 1.4 ± 0.6 | 2.5 ± 1/0 |
| $\gamma_2$ (%) | 44 ± 3.0 | 63 ± 5.2 | 72 ± 24 | 87 ± 12 | 34 ± 4.0 |
| Reduced $\chi^2$ | 2.4 | 3.0 | 0.8 | 0.3 | 3.2 |
| PTC as a function of conversion | | | | | |
| Cellulose | SC | BC | FP | MCC | AC |
| $S_{P1}$ (µmole/g cellulose) | 5.0 ± 1 | 4.0 ± 0.5 | 2.0 ± 0.6 | 1.6 ± 0.1 | 1.4 ± 0.3 |
| $S_{P2}$ (µmole/g cellulose) | 0.7 | 0.7 ± 0.5 | 1.6 ± 0.2 | 0.7 ± 0.1 | 0.6 ± 0.3 |
| $\gamma_1$ (%) | 1.7 | 2.5 ± 6.2 | 0.3 ± 0.03 | 0.2 ± 0.003 | 7.0 ± 0.2 |
| $\gamma_2$ (%) | 65 | 116 ± 10 | 57 ± 4.8 | 71 ± 3.1 | 185 ± 8.8 |
| Reduced $\chi^2$ | 1.8 | 6.7 | 0.7 | 0.3 | 4.4 |

### Bound enzyme populations for swollen cellulose, filter paper, and microcrystalline cellulose

The decrease in overall hydrolysis rates and extents of conversion of BC (Figure 8A) was accompanied by a drop in the maximum concentration of productively bound enzyme on BC from 0.02 to ~0.001 µmole/g even though the concentration of productive binding sites remained unchanged (Figure S7C). The increased concentration of non-productive binding sites did result in increased concentrations of non-productively bound enzyme (Figure S7A). Non-productive binding decreased with the lower concentration of non-productive binding sites on AC (Figure S7B). Overall, we observed that maintaining a higher ratio of productive to non-productive binding sites favors productive binding and thus increases hydrolysis rates.

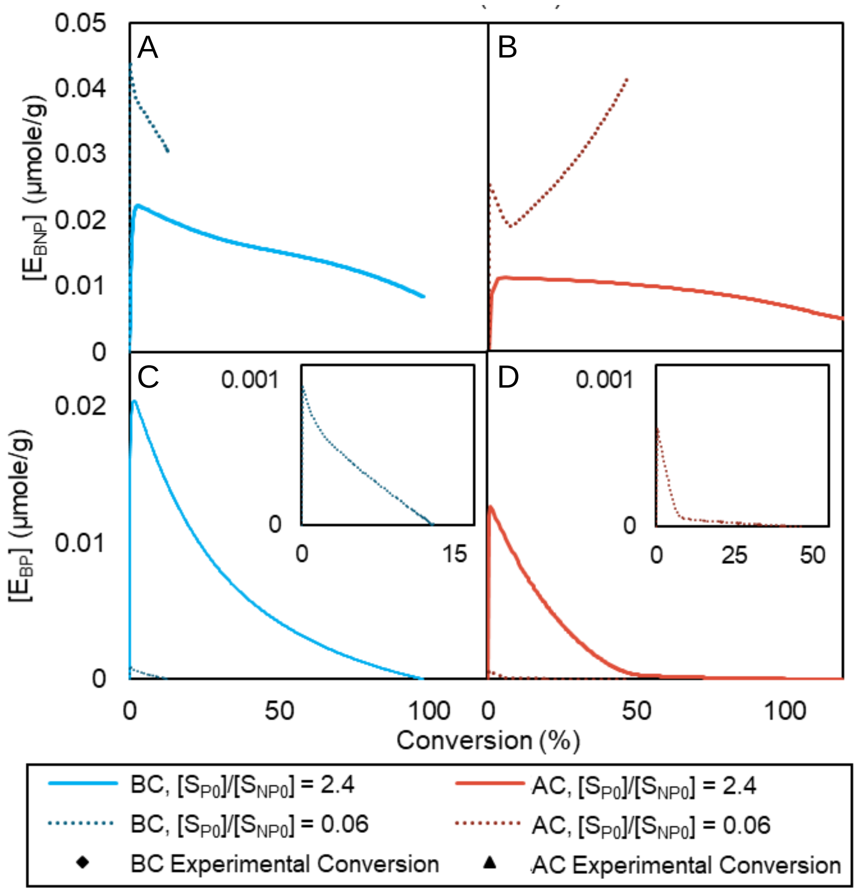


*Figure S7: A) Simulated conversion of BC with $[S_{P0}]/[S_{NP0}]$ ratio of 2.4 and 0.06 over time (alongside experimental conversion of BC from Figure 5A (main text); B)simulated conversion of AC with $[S_{P0}]/[S_{NP0}]$ ratio of 2.4 and 0.06 with respect to time (alongside experimental conversion of AC from Figure 5B (main text); model predictions for the concentration of non-productively bound enzyme with respect to conversion of BC (C) and AC (D) with $[S_{P0}]/[S_{NP0}]$ ratios of 2.4 and 0.06; model predictions of the concentration of productively bound enzyme with respect to conversion of BC (E) and BC (F) with $[S_{P0}]/[S_{NP0}]$ ratios of 2.4 and 0.06; inset in (F) is a zoomed plot of the concentration of non-productively bound enzyme on AC with $[S_{P0}]/[S_{NP0}]$=0.06.*

An accumulation of non-productively bound enzymes at high conversions was observed on AC (Figure S7B) as well as on the other recalcitrant celluloses, FP and MCC (Figure S8). The decrease in the $[S_{P0}]/[S_{NP0}]$ ratio of BC to 0.06 by increasing non-productive binding site concentrations increased non-productive binding, but did not result in the accumulation of non-productively bound enzyme (Figure S7A). However, when we increased the productive to non-productive binding site ratio of AC to 2.4 by decreasing non-productive binding site concentrations, the magnitude of non-productive binding decreased about two-fold, and the buildup of non-productively bound enzymes was eliminated (Figure S7B). This suggests that elevation of the initial productive binding site fraction by eliminating non-productive binding sites (e.g., through pretreatment) can increase burst phase hydrolysis rates, but preventing non-productive binding by retaining a higher fraction of productive binding sites throughout conversion (e.g., by synergistic enzyme action with accessory enzyme) is also key to overcoming hydrolysis limitations.

### Bound enzyme populations for swollen cellulose, filter paper, and microcrystalline cellulose

The model predicted populations of productively and non-productively bound enzyme are shown in Figure S8 for swollen cellulose, filter paper, and microcrystalline cellulose (bacterial and algal cellulose are shown in Figure S7).

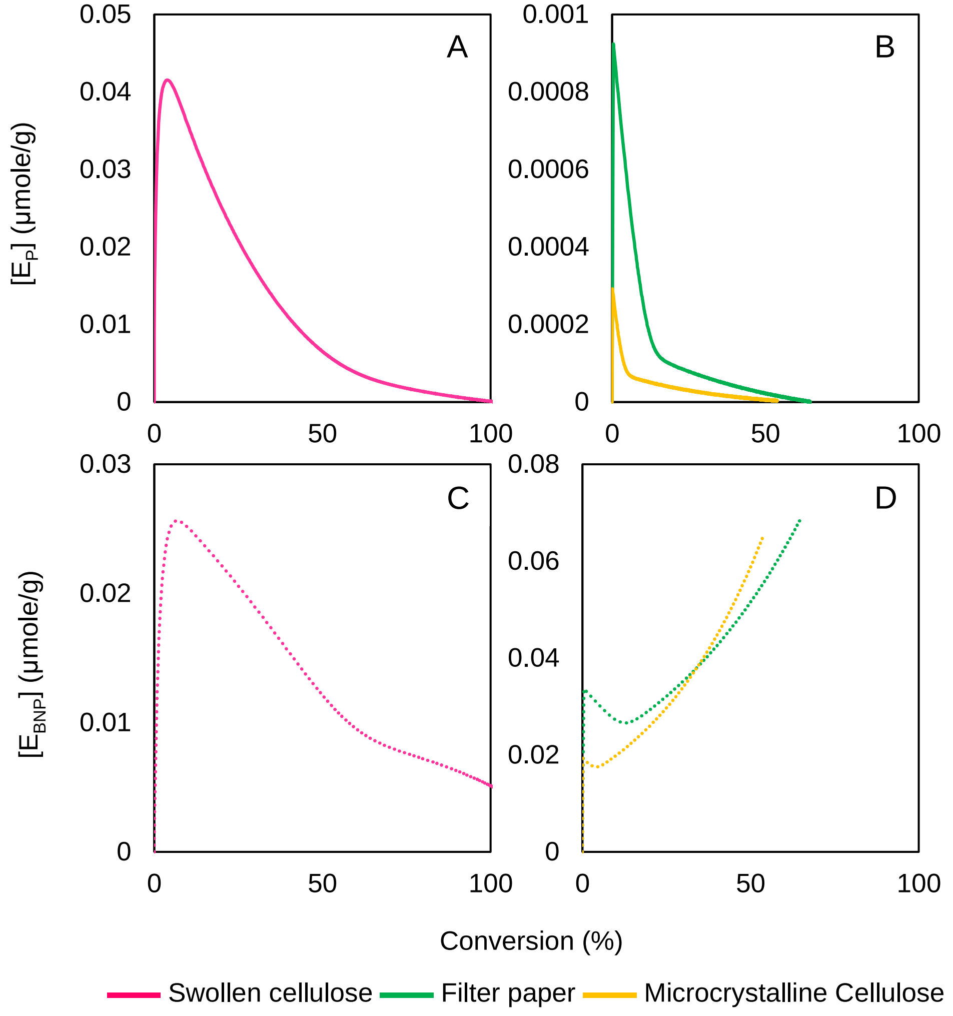


*Figure S8: Simulated bound enzyme concentrations (µmole/g) (using the model described in Table 1 in the main text). The concentration of productively bound enzyme on A) swollen cellulose (pink); B) filter paper (green) and microcrystalline cellulose (yellow); Non-productively bound enzyme concentration on C) swollen cellulose and D) filter paper and microcrystalline cellulose.*

### Complexed, but stalled non-productive enzyme binding shortens burst phase hydrolysis

Since the abundance of non-productive binding sites appears to significantly impact hydrolysis rates, we also examined the role of non-productive enzyme binding on hydrolysis rates. We simulated scenarios where non-productive binding steps were systematically eliminated (Figure S9A). Eliminating non-productive binding from solution (Scenarios 1, 2 and

3) had little effect on hydrolysis rates of either the easily digestible BC or the recalcitrant AC (Figure S9B and C). On BC, eliminating non-productive complexation from solution (Scenario 1) resulted in a faster decrease in the concentration of non-productively bound enzyme (Figure S9D), while on AC, an overall decrease in non-productive binding was observed and non-productively bound enzyme no longer accumulated on the substrate (Figure S9E). Productive binding was not impacted for either substrate (Figure S9F and G). Eliminating the formation of non-productively bound enzyme by decomplexation of productively bound enzyme (Scenario 3) caused a slight increase in hydrolysis rates (Figure S9B, C) after the initial burst phase. Overall, decreasing long term non-productive enzyme buildup by either eliminating non-productive binding from solution or eliminating decomplexation of a surface-bound enzyme (Scenarios 1, 2, or 3) had negligible impact on productive binding and conversion (Figure S9).

Eliminating non-productive enzyme binding due to stalling of complexed enzymes (Scenario 4) greatly increased overall conversion by extending the burst phase (Figure S9B and C). Without enzyme stalling, the accumulation of non-productive enzyme on AC persisted but lessened, while non-productive binding decreased on BC. Moreover, productive binding increased significantly on both BC and AC (Figure S9F and G). A similar extension of the burst phase was observed in simulations where irreversibly bound enzyme was removed to expose blocked productive binding sites (Figure 8), suggesting that enzyme stalling upon encountering a non-productive binding site is the origin of these irreversibly bound enzymes. As our method does not distinguish between enzymes bound with occupied or free active site, we cannot comment on the binding mode of enzymes remaining on the cellulose surface after salt washing. However, evidence in the literature points to buildup of processive enzymes upon encountering the end of an “obstacle free path” on the substrate as a source of strongly bound enzymes[16-23]. Decreasing hydrolysis rates have been observed while the concentration of either the total adsorbed cellulase[24] or Cel7A bound by the active site remained constant (at < 10 hours)[20]. Taken with the simulation results, we thus conclude that the persistence of the burst phase is limited by non-productive binding when complexed enzymes stall (Step 5 in Figure 2).

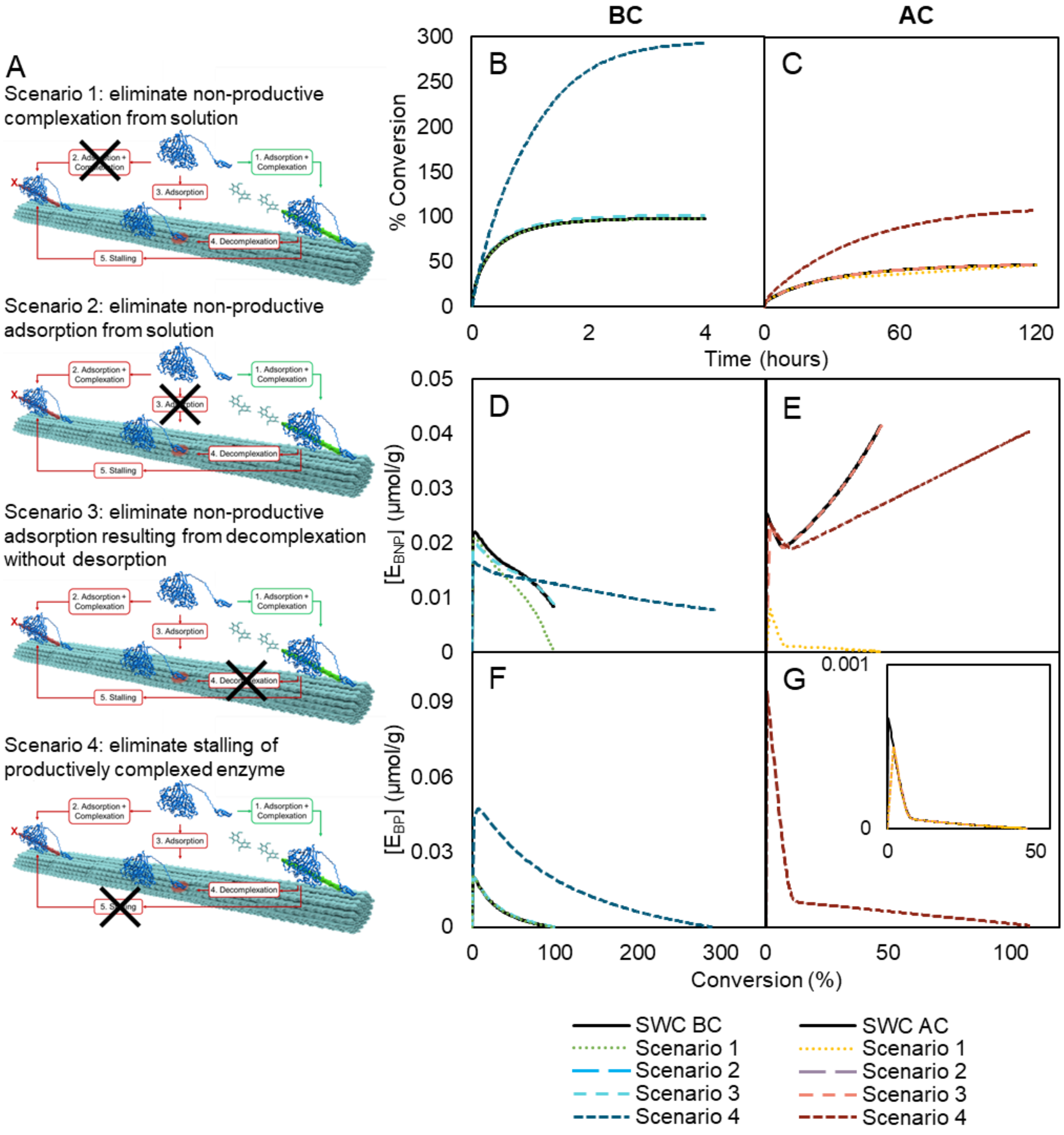


*Figure S9: A) Simulations were conducted with removal of binding scenarios from Figure 2 (main text). Simulations of scenarios 1-4 as shown in A): B) Simulated conversion of BC; C) Simulated conversion of AC; D) Non-productively bound enzyme on BC; E) Non-productively bound enzyme on AC; F) Productively bound enzyme on BC; G) Productively bound enzyme on AC.*
